# Unbiased mapping of cereblon neosubstrate landscape by high-throughput proteomics

**DOI:** 10.1101/2024.10.18.618633

**Authors:** Martin Steger, Gisele Nishiguchi, Qiong Wu, Bjoern Schwalb, Bachuki Shashikadze, Kevin McGowan, Zhe Shi, Jeanine Price, Anand Mayasundari, Lei Yang, Anastasia H. Bednarz, Sophie Machata, Tobias Graef, Denis Bartoschek, Vadim Demichev, Uli Ohmayer, Jun Yang, Henrik Daub, Zoran Rankovic

## Abstract

Molecular glue degraders (MGDs) are small molecules that harness the ubiquitin-proteasome system to induce degradation of target proteins, including those lacking conventional ‘druggable’ pockets. Given the challenges in their rational design, MGD discovery predominantly relies on screening-based approaches, such as cell viability assays. However, one potential limitation of such screening methods is the risk of overlooking non-essential neosubstrates of potential therapeutic value. To address this concern, we present a high-throughput proteome-wide MGD screening platform utilizing label-free, data-independent acquisition mass spectrometry (DIA-MS) for integrated proteomics and ubiquitinomics analysis. Processing a diverse set of 100 CRBN-ligands across two cancer cell lines reveals a broad array of neosubstrates, including 50 novel candidates validated by MS-based ubiquitinomics. These findings considerably expand the current landscape of CRBN-mediated neosubstrates. Comprehensive hit validation and structure-degradation relationship analyses guided by global proteomics, identifies highly selective and potent phenyl glutarimide-based degraders of novel neosubstrates, including KDM4B, G3BP2 and VCL, none of which contain the classical CRBN degron motif. This study demonstrates that comprehensive, high-throughput proteomic screening offers new opportunities in MGD drug discovery.

## Introduction

Targeted protein degradation (TPD) is a rapidly emerging drug discovery paradigm, which relies on engagement of normal cellular proteolytic processes to degrade disease-associated proteins^1^. The two currently most advanced TPD approaches employ proteolysis targeting chimeras (PROTACs)^2^ or molecular glue degraders (MGDs)^3^. MGDs are small molecules that can bind and alter specificity of an E3 ligase, resulting in recruitment, ubiquitination, and subsequent degradation of new, non-physiological substrates, commonly referred to as neosubstrates. The immunomodulatory drugs (IMiDs) thalidomide, lenalidomide and pomalidomide, clinically used as the first-line therapy for multiple myeloma, were the first drugs found to exert their therapeutic effect through such a mechanism^4^. IMiDs bind to cereblon (CRBN), a substrate recognition subunit of CRL4^CRBN^ ligase^5^, and induce degradation of the transcription factors IKZF1 and IKZF3 that are required for multiple myeloma cell survival^6,7^. Importantly, the neosubstrate recruitment is governed by MGD binding to the E3 ligase creating a new, neomorphic recognition surface and does not require a ligandable pocket^8,9^. This mechanism provides an unprecedented and exciting opportunity to tackle currently undruggable therapeutic targets, such as transcription factors.

Interestingly, despite their high structural similarity, lenalidomide and pomalidomide display different protein degradation profiles. While both degrade the transcription factors IKZF1/3, only lenalidomide induces degradation of CSNK1A1 (CK1α), indicating how a small difference in degrader molecular structure can significantly alter the neosubstrate specificity. Indeed, an ever-increasing number of neosubstrates have been discovered in recent years with each IMiD displaying distinct degradation patterns, supporting the notion that E3 ligase substrate specificity can be modulated by structural alterations of the ligand. Unfortunately, prospectively designing such molecules and rationalizing the structure degradation relationship remains to be highly challenging.

Mass spectrometry (MS)-based proteomics has undergone major advancements in recent years, driven by deep learning-enhanced software and next-generation MS hardware^10–12^. These developments enable near-complete and highly reproducible proteome quantifications in high-throughput mode. A key advantage of MS-based proteomics is its unbiased, target- and E3-ligase agnostic screening capability, allowing for the simultaneous discovery of novel targets and evaluation of degrader selectivity. Despite its routine application in profiling selected degrader candidates, integrated, high-throughput screening workflows for MGDs remain underdeveloped.

We recently synthesized a chemically diverse library of currently over 5,000 CRBN ligands and screened it against a large panel of cancer cell lines in viability assays, aiming at discovering novel degraders and cancer vulnerabilities. This proved to be a productive effort that delivered potent and selective degraders of proteins such as G1-to-S-phase transition (GSPT1)^13,14^ and casein kinase 1A (CK1α)^15,16^ that exhibited antiproliferative activity against a broad range of human cancer cell lines. However, despite these encouraging results, our phenotypic screening approach relied solely on the cell viability as the assay readout. As a result, proteins that are not essential for survival of cancer cells in our screening panel would be missed by such an approach. To investigate the potential extent of such omissions and further expand the CRBN neosubstrate landscape, we selected a representative subset of our tractable molecular glue library for in-depth characterization by MS-based proteomics.

## Results and Discussion

### Deep proteomic screening of a molecular glue degrader library

Our IMiD-based molecular glue library was designed and synthesized based on the previously described principles and protocols. To expand the chemical diversity of our library beyond IMiDs, we included multiple CRBN-binding cores, including those recently reported, such as phenyl-glutarimide (PG) and several proprietary (other) scaffolds^17^. For each chemical core, we scanned vectors with different functionalities, applied an array of chemistries and used a diverse set of building blocks to construct compounds covering a broad range of chemical space and physicochemical properties. The compound selection criteria for mapping the neosubstrate landscape using proteomics was defined based on scaffold diversity and we selected one representative compound based on the average molecular weight within the scaffold and powder availability. We excluded scaffolds known to degrade GSPT1 or CK1α based on our phenotypic screening campaign but included our previously reported CK1α (SJ10040) and GSPT1 (SJ6986) degraders as controls. In this fashion, we selected 57 compounds belonging to the IMiD scaffold (35 lenalidomide and 22 thalidomide core analogues), 41 PGs and 2 compounds from other CRBN binding scaffolds (Supplementary Fig.1). We next designed a multilayered proteomics workflow to screen the 100 selected MGDs in both Huh-7 and in NB-4 cells, aiming to expand the number of targetable proteins through CRBN-based degraders and gain deeper insights into their mechanisms of action. We selected these two cell lines based on their complementary protein expression profiles, allowing us to maximize the number of targetable proteins for our degraders. Our primary screen consisted of 24-hour compound treatments at 10 µM in 96-well plates, followed by semi-automated proteomics sample preparation and label-free, data-independent acquisition mass spectrometry (diaPASEF)^18^, enabling deep proteomics screening with high-throughput. In total we profiled one hundred compounds in triplicate treatments in both cell lines, along with 120 DMSO controls, resulting in 720 LC-MS runs. Following MS data processing and statistical analysis, we identified proteins with a reduction in abundance of over 25% as candidate CRBN neosubstrates and assessed their degradation kinetics at 6 hours of MGD treatment. For candidates significantly downregulated at both time points, we confirmed cullin-RING E3 ligase (CRL)-dependency by co-treating cells with the respective MGD and the NEDD8 E1 inhibitor MLN4924^19^. Finally, to mechanistically validate those putative neosubstrates, we performed global ubiquitinomics. Proteins that were rapidly ubiquitinated in cells following MGD treatment were considered bona fide CRBN neosubstrates (Fig. 1a and Supplementary Table 1).

**Figure 1.**
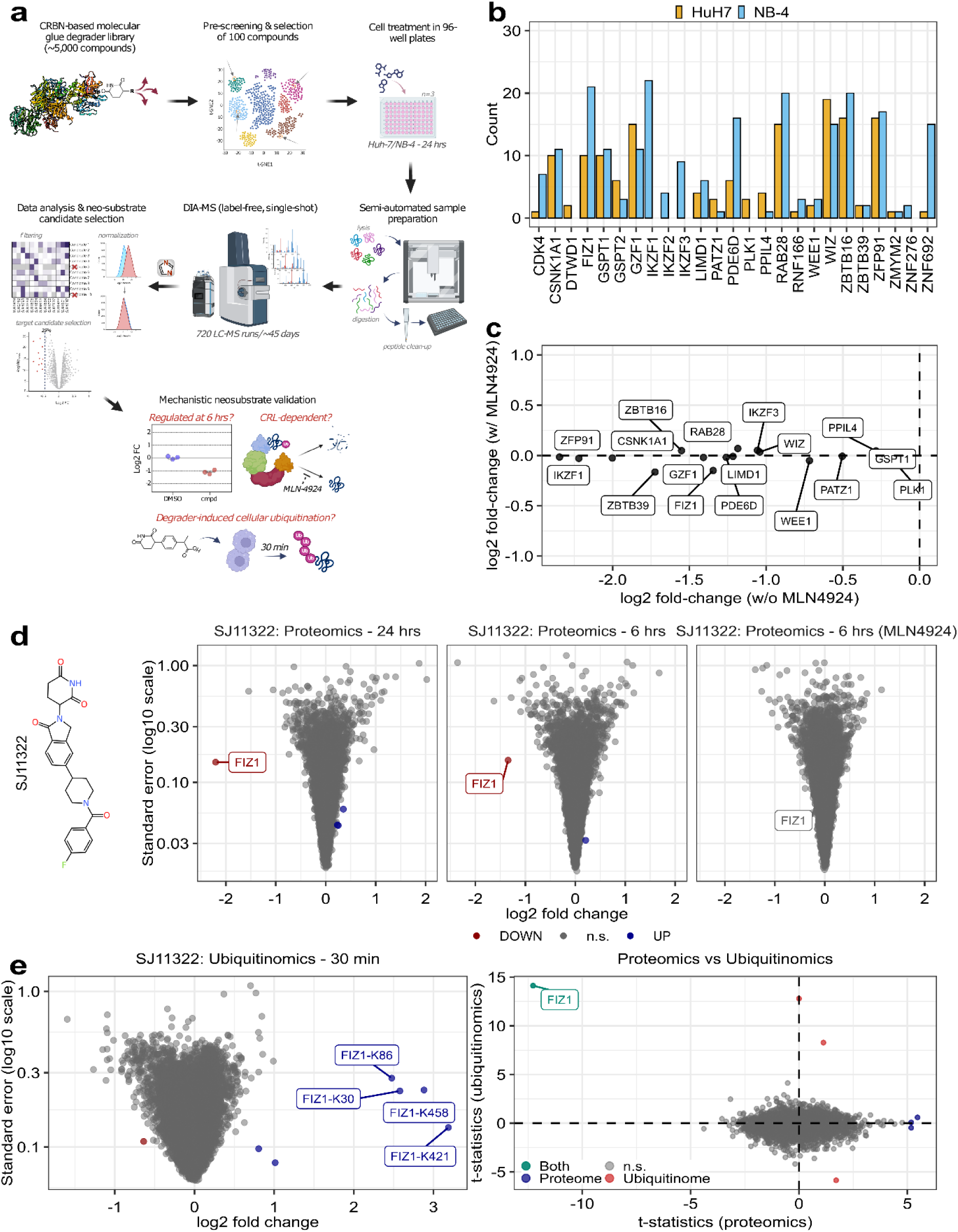
The molecular glue degrader (MGD) proteomics screening workflow. (A) Schematic of the proteomics screening workflow. A subset of 100 CRBN-based molecular glue degraders (MGDs) was selected from a library of 5,000 compounds and used for cell treatments at a concentration of 10 µM in Huh-7 and NB-4 cells, plated in 96-well plates. Each plate included 25 MGDs tested in triplicate. Following sample preparation, mass spectrometry (MS) data were acquired using a Bruker TimsTOF platform, and the raw data was processed with DIA-NN^10^ software. After filtering for missing values and normalizing the data, proteins that were significantly downregulated by more than 25% were selected for re-testing using a 6-hour compound treatment, either in the absence or presence of the protein neddylation inhibitor MLN4924. Ubiquitinomics analysis was subsequently employed to mechanistically validate MLN4924-dependent neosubstrate candidates. (B) Count of significant downregulation events (LIMMA^29^, FDR=1%) detected in Huh-7 or NB-4 cells for previously reported neosubstrates (see also Supplmentary Table 2). The compound treatment time was 24 hours. (C) A scatter plot of known CRBN neosubstrates identified in Huh-7 or NB-4 cells, showing downregulation upon MGD treatment (6 hours), which was suppressed following co-treatment with MLN4924 (5 µM). (D) Volcano plots depicting protein quantifications for the selective FIZ1 degrader SJ11322 (10 µM), with log_2_-transformed fold-change on the x-axis and standard error (log_10_ scale) on the y-axis. The plots, from left to right, represent: (i) 24-hour treatment, (ii) 6-hour treatment, and (iii) 6-hour treatment with SJ11322 in the presence of MLN-4924 (5 µM). The chemical structure of SJ11322 is shown on the left side. Significant up- and downregulations (proteins (LIMMA^29^, FDR=1%) or ubiquitination sites (LIMMA^29^, FDR=5%)) are colored in blue and red, respectively. n.s.= not significant. (E) Volcano plots displaying ubiquitinated peptide quantifications for SJ11322. The treatment time was 30 minutes and significantly upregulated ubiquitinated peptides are colored in blue (LIMMA^29^, FDR=5%). Four FIZ1 ubiquitination sites that were induced upon compound treatment are highlighted. (E) Comparison plot of the t-statistics (SJ11322 vs. DMSO) of the proteome (x-axis) and the ubiquitinome (y-axis). The t-statistics of significantly upregulated FIZ1 ubiquitination sites were averaged. Statistically significant up- and downregulations in the proteome (LIMMA^29^, FDR=1%) and ubiquitinome (LIMMA^29^, FDR=5%) are colored in blue and red, respectively. Proteins that were significantly downregulated after 24 hours of compound treatment and exhibited upregulated ubiquitination sites at 30 minutes of compound treatment are highlighted in turquoise. n.s.= not significant.

Our proteomic screening campaign quantified an average of 10,200 protein groups per compound in Huh-7 and 9,500 in NB-4 cells, with over 98% of these retained for statistical testing after filtering for missing values. The median coefficient of variation (CV) for protein groups (n = 3) was about 6% in both cell lines (Supplementary Fig. 2a,b). This highlights that our single-shot MS-based screening platform detects a large proportion of the proteome expressed by a human cell line, with very high data completeness and excellent quantification precision^11,20^.

We first classified the compounds based on the number of significantly induced protein downregulations. About 25% of the compounds did not modulate any proteins in either cell line, while approximately 50% induced specific changes, with 10 or fewer proteins displaying a significant reduction in abundance. The remaining compounds were more promiscuous, inducing numerous protein regulations, some of which may reflect secondary effects resulting from a prolonged drug exposure. Due to their modulation of thousands of proteins, SJ6986 and SJ41478 were excluded from neosubstrate identification analyses (Supplementary Fig. 2c).

We identified 35 neosubstrates for which a decrease in cellular abundance following treatment with CRBN-directed degrader molecules has been previously demonstrated^21–24^. 33 of these were quantified in either Huh-7 or NB-4 cells and 25 were significantly downregulated after 24 hours of treatment by at least one compound (Supplementary Table 2). Among those, RAB28, FIZ1, ZBTB16 and ZFP91 emerged as the most frequent hits common to both cell lines. In contrast, Ikaros family zinc finger proteins IKZF1, IKZF2 and IKZF3 were exclusively detected and downregulated in NB-4 cells, consistent with their immune cell-specific expression (Fig. 1b)^25^. Notably, compared to Huh-7, strong IKZF1 degraders consistently induced a greater number of protein regulations in NB-4, likely due to secondary effects resulting from inhibition of IKZF1-mediated downstream signaling (Supplementary Fig. 2d).

To assess degradation kinetics, we performed 6-hour treatment experiments to re-test the compounds that induced the strongest effects on each neosubstrate following 24 hours of drug exposure. Many neosubstrates displayed comparable levels of downregulation at both time points; however, proteins like FIZ1 and LIMD1 showed a more pronounced downregulation at the 24-hour time point (Figure 1c and Supplementary Fig. 3). In line with a CRL-dependent mechanism, a 6-hour co-treatment of MGDs with MLN4924 reversed the downregulation of all neosubstrates (Fig. 1c). Notably, our screening efforts uncovered selective degraders of previously reported IMiD neosubstrates, such as LIMD1, GZF1, and FIZ1. (Fig. 1d and Supplementary Fig. 3)^21,23,26^. Since intracellular ubiquitination of these proteins at endogenous levels following treatment with CRBN-based degraders has not yet been demonstrated, we subsequently conducted a series of global ubiquitinomics analyses (Supplementary Fig. 4)^27,28^. We reasoned that rapid cellular ubiquitination of target proteins upon drug exposure is driven exclusively by MGD-induced proximity between CRL4^CRBN^ E3 ligase and its neosubstrate and consequently we selected a 30-minute treatment time. As a proof-of-concept, we globally profiled protein ubiquitination in response to the newly identified and selective FIZ1 degrader SJ11322. This revealed several strongly upregulated ubiquitination sites on FIZ1, providing direct evidence of its ubiquitination following degrader treatment, and further corroborating previous findings that FIZ1 is a neosubstrate of CRBN^26^ (Fig. 1e). Collectively, these results highlight that the combination of our MGD library design with MS-based proteomics and ubiquitinomics can identify and validate high-quality degrader probes for poorly characterized neosubstrates.

### Systematic discovery of novel CRBN neosubstrates

We next analyzed our proteomics screening data for putative novel CRBN neosubstrates, initially focusing on the Huh-7 cells. Out of the 100 compounds tested at 24 hours, 32 resulted in a decrease in cellular abundance of at least one protein by more than 25%. This led to a preliminary list of 84 novel neosubstrate candidates. The 32 hit compounds were retested at the 6-hour time point, resulting in the identification of 16 compounds that downregulated 26 potentially novel neosubstrates. Although the downregulation of UHRF1, ZNF430, ZNF644, DHX37 and CCNK in the 24-hour dataset was slightly below statistical significance or the 25% fold-change cut-off, these proteins exhibited significant and robust downregulation following the 6-hour treatment. We therefore included them in the follow-up experiments (Supplementary Table 3). For each of the neosubstrate candidates, we selected at least one compound and performed co-treatment experiments with MLN4924. This strongly inhibited the downregulation of 23 proteins, displaying more than a 70% reduction compared to MGD treatment alone, thereby classifying them as putative novel CRBN neosubstrates (Fig. 2a). We also recorded CRL-independent downregulation events, including those of the cytochrome P450 enzyme CYP2J2 and oxysterol-binding protein-related protein 8 (OSBPL8), likely reflecting CRBN-independent off-target effects (Supplementary Fig. 5). We next subjected the 23 CRL-dependent neosubstrate candidates to quantitative ubiquitinomics. These experiments revealed a total of 16 novel neosubstrates, including the lysine-specific histone demethylase 4B (KDM4B)^30^, the CCR4-NOT transcription complex subunit 4 (CNOT4)^31^, the zinc finger protein ZBTB7B^32^, and the multifunctional protein folliculin (FLCN)^33^ (Fig. 2b,c, Supplementary Fig. 6 and Supplementary Table 3). We also recorded ubiquitination and subsequent degradation of several known protein complexes, such as CSNK1A1 and its interaction partners FAM83G and FAM83B, following treatment with SJ10040 (Supplementary Fig. 7a)^34^. The recruitment of FAM83G or FAM83B by CSNK1A1 likely positions them in proximity to the MGD-bound CRBN, thereby facilitating their ubiquitination as bystander neosubstrates^35–37^. Additional examples of potential co-degradation included CDK-activating kinase (CAK) complex members CDK7, CCNH and MNAT1 (SJ42101), as well as CDK13 and its interaction partner CCNK (SJ42509)^38,39^ (Supplementary Fig. 7b,c). In case of the CAK complex, MNAT1 appears to be the direct MGD target, as suggested by its stronger downregulation and ubiquitination. This is further supported by recent findings demonstrating MGD-induced in vitro and cellular recruitment of MNAT1 to CRBN via a β-hairpin G-loop recognition motif, which is absent in both CDK7 and CCNH^36,40^. In contrast to previous studies that identified potent CCNK degraders with weaker co-degradation effects on CDK12 and CDK13, the MGD SJ42509 induced only modest CCNK degradation while exhibiting a stronger effect on CDK13. This evidence, along with the presence of a G-loop motif, suggests that CDK13 is a direct target of SJ42509. Furthermore, it indicates a distinct mode of action compared to previously reported MGDs, which deplete CCNK through a drug-induced interaction of CDK12 or CDK13 with DDB1^41–43^.

**Figure 2.**
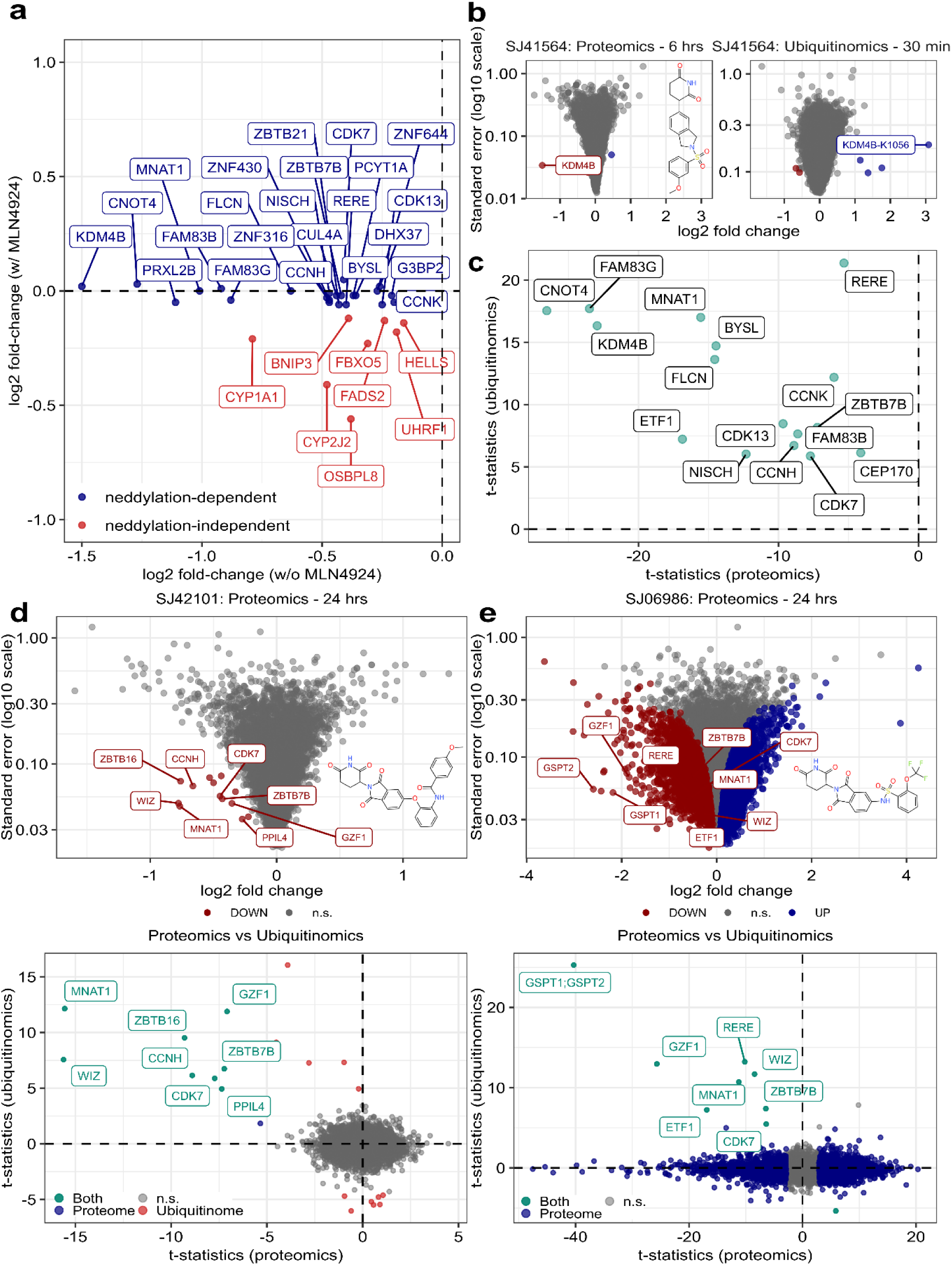
Discovery of novel neosubstrates in Huh-7 cells. (A) Scatter plot illustrating newly identified CRBN neosubstrates in Huh-7 cells. The plot displays all proteins that showed significant downregulation following treatment with 10 µM MGD for 6 hours (x-axis) alongside their corresponding fold-changes under co-treatment with 5 µM MLN4924 (y-axis). Proteins regulated in a neddylation-dependent manner are highlighted in blue, while those regulated in a neddylation-independent manner are indicated in red. (B) Volcano plots depicting protein (left) and ubiquitinated peptide (right) quantifications for the KDM4B degrader SJ41564 (10 µM), with log_2_ fold-change on the x-axis and standard error on the y-axis (log_10_ scale). Significant up- and downregulations of proteins (LIMMA^29^, FDR=1%) or ubiquitination sites (LIMMA^29^, FDR=5%) are colored in blue and red, respectively. (C) Summary of proteome (x-axis) and ubiquitinome (y-axis) t-statistic comparisons for all newly validated neosubstrates in Huh-7 cells. T-statistics for ubiquitination sites mapped to the same protein were averaged, and only significantly upregulated sites were included in the analysis. (D) and (E) Volcano plots of protein quantifications for SJ42101 (D) and SJ06986 (E) after 24 hours of treatment. Significantly up- and downregulated proteins (LIMMA^29^, FDR=1%) are colored in blue and red, respectively. The proteome (x-axis) and ubiquitinome (y-axis) t-statistic comparison plots are shown (bottom). Statistically significant up- and downregulations in the proteome (LIMMA^29^, FDR=1%) and the ubiquitinome (LIMMA^29^, FDR=5%) are colored in blue and red, respectively. Proteins that are both significantly downregulated after a 24-hour compound treatment and harbor upregulated ubiquitination sites at 30 minutes of compound treatment are colored in turquoise. n.s.= not significant.

We identified two compounds (SJ42101 and SJ6986) that downregulated MNAT1. Treatment of cells for 24 hours with SJ42101 yielded only 13 downregulated proteins. In contrast, the potent GSPT1/GSPT2 degrader SJ6986, at a concentration of 10 µM and the same time point, modulated the abundances of thousands of proteins, obscuring direct protein degradation events through protein modulations resulting from disrupted protein biosynthesis^14,44^. To distinguish between direct target degradation and indirect GSPT1-mediated effects, we conducted global ubiquitinomics with both compounds. While SJ42101 induced ubiquitination of 7 neosubstrates, treatment with SJ6986 resulted in the ubiquitination of GSPT1, GSPT2 and their complex partner ETF1^45^, as well as the proteins MNAT1, CDK7, RERE, GZF1, WIZ and ZBTB7B (Fig. 2d,e). This demonstrates that our integrated proteomics and ubiquitinomics screening approach enables the specific extraction of primary degrader targets from complex proteomic profiles, such as those induced by GSPT1 degradation.

To assess potential cell line-specific responses of MGDs, we extended our proteomic screening efforts to NB-4 cells. While we could recapitulate most CRL-dependent protein downregulations identified in Huh-7 (e.g. KDM4B or CNOT4), some effects were cell line-specific. For instance, ZBTB7B was downregulated by SJ42101 only in Huh-7, while SJ10040 affected the ubiquitous protein kinase CSNK1E exclusively in NB-4 cells (Supplementary Fig. 8a,b). Similarly, both SJ42101 and the KDM4B degrader SJ41564 significantly downregulated vinculin (VCL) in NB-4, but not Huh-7, despite its ubiquitination in both cell lines (Supplementary Fig. 8c). These results demonstrate that MGDs can induce cell type-specific regulations even for ubiquitously expressed proteins. While the underlying mechanisms are not yet fully understood, identifying novel cell type-specific CRBN neosubstrates underscores the benefit of screening compounds across different cell lines.

### Overexpression of CRBN enhances neosubstrate identification

Our ubiquitinomics-based validation in Huh-7 cells identified 16 novel neosubstrates (Figure 2c) and we further validated VCL as CRBN neosubstrate in NB-4 cells. However, we were unable to detect compound-induced ubiquitination for the remaining 11 neosubstrate candidates. These were ZBTB21, CSNK1E, ZNF316, CUL4A, ZNF644, PRXL2B, G3BP2, ZNF430, DHX37, PCYT1A and STK38 that were regulated by at least one out of eight MGDs identified in the proteomic screens (Supplementary Table 4). These target candidates were generally expressed at low levels or only modestly downregulated by the tested degraders, indicating potential sensitivity limitations despite the quantification of up to 30,000 ubiquitination sites. We hypothesized that CRBN overexpression would enhance cellular CRL4^CRBN^ activity to both increase ubiquitination and subsequent degradation of neosubstrates, as previously demonstrated for lenalidomide targets^46^. We therefore generated a HEK293 cell line with stable CRBN overexpression (HEK293-CRBNoe), which exhibited about eight-fold higher levels of CRBN compared to the parental cells (Supplementary Fig. 9). To verify whether increased CRBN protein expression was indeed associated with enhanced degradation efficiency, we first compared the proteomic profiles of SJ10040-treated HEK293 and HEK293-CRBNoe cells. As anticipated, a 6-hour compound treatment of HEK293-CRBNoe cells led to increased degradation of the known neosubstrates CSNK1A1, WIZ and WEE1 (Fig. 3a). The putative neosubstrates ZBTB21 and CSNK1E, initially identified in the Huh-7 and NB-4 screens, were significantly downregulated exclusively in HEK293-CRBNoe cells. In addition to confirming neosubstrate candidates from the primary screen, we discovered novel putative neosubstrates that were exclusive to HEK293-CRBNoe, such as ZNF324, ZBTB41, STK35 and CHD7. For example, ubiquitinomics analysis of SJ10040 revealed rapid intracellular ubiquitination of 13 neosubstrates that included both ZBTB21 and CSNK1E, as well as the four newly identified target candidates, validating them as CRBN neosubstrates (Fig. 3b). Proteomics and ubiquitinomics analysis of the remaining seven compounds in HEK293-CRBNoe cells confirmed their enhanced response to MGDs, further validating CUL4A, ZNF316, PRXL2B, G3BP2, ZNF644, ZNF430, DHX37 and PCYT1A as novel neosubstrates (Supplementary Fig. 10,11). Interestingly, global ubiquitination profiling of the promiscuous degrader compound SJ42101 in HEK293-CRBNoe cells uncovered about 30 neosubstrates, including many novel ones, such as the zinc finger proteins ZNF121, ZNF239, ZNF408 and ZNF507, the peptidase ZUP1, and the mRNA 3’-end processing factor CSFT2T (Fig. 3b). Collectively, our integrated proteomics and ubiquitinomics screening strategy revealed 50 novel neosubstrates from a mere selection of 100 compounds, demonstrating a much broader target scope for CRBN-based degraders than previously recognized (Fig. 3c, Supplementary Fig.12 and Supplementary Table 5).

**Figure 3.**
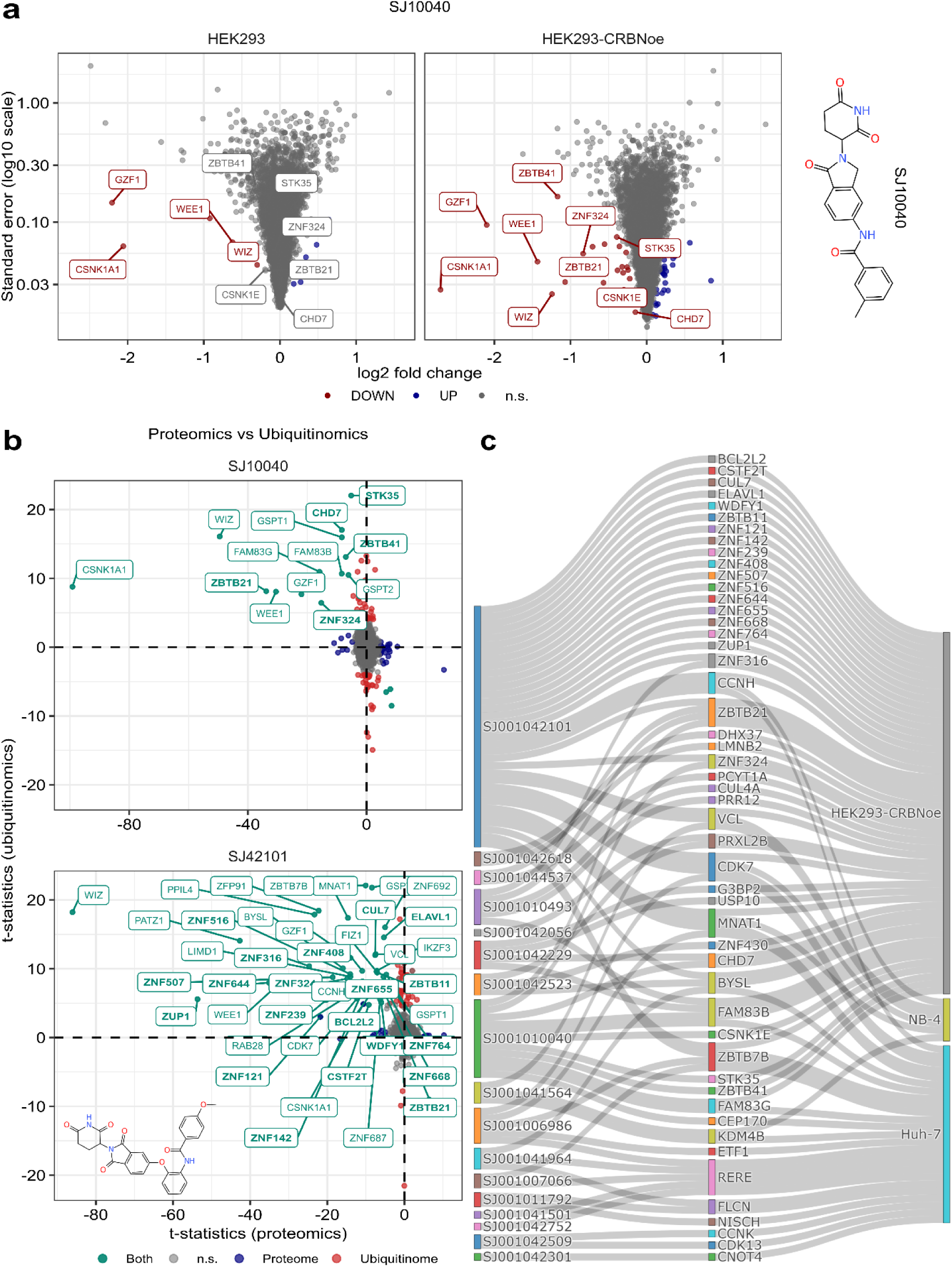
Neosubstrate discovery in HEK293-CRBNoe cells. (A) Volcano plots showing protein quantifications of HEK293 (left) and HEK293-CRBNoe (right) in response to the degrader compound SJ10040. The x-axis represents log_2_ fold change, while the y-axis shows standard error on a log_10_ scale. Both cell lines were treated with the compound for 6 hours. Significantly up- and downregulated proteins, as determined by LIMMA^29^ (FDR = 1%), are indicated in blue and red, respectively. The chemical structure of SJ10040 is shown on the right side. n.s.= not significant. (B) Comparison plot of the t-statistics of the proteome (x-axis) and the ubiquitinome (y-axis) for the two MGDs SJ10040 (top) and SJ42101 (bottom) in HEK293-CRBNoe cells. The t-statistics of significantly upregulated ubiquitination sites mapping to the same gene were averaged. Statistically significant up- and downregulations of the proteome (LIMMA^29^, FDR=1%) and the ubiquitinome (LIMMA^29^, FDR=5%) are colored in blue and red, respectively. Proteins that were significantly downregulated after 6 hours of compound treatment and exhibited upregulated ubiquitination sites at 30 minutes of compound treatment are highlighted in turquoise. Neosubstrates that were identified and validated in HEK293-CRBNoe are indicated in bold. n.s.= not significant. (C) Sankey diagram summarizing all novel neosubstrates that were identified and validated through ubiquitinomics in Huh-7, NB-4 and HEK293-CRBNoe cells. The compounds are displayed on the left side of the diagram, while the cell lines are represented on the right side.

### Impact of MGD structure on degradation specificity and selectivity: Structure-degradation relationship

Almost all screened compounds (98 out of 100) either contained classical IMiD core structures as present in thalidomide and lenalidomide or possessed a phenyl-glutarimide (PG) core recently introduced as alternative CRBN-binding warhead^47^. Our comprehensive proteomics data for these compounds allowed us to explore correlations between MGD chemistry and cellular degradation activities. To do that, we first grouped all compounds that degraded known or newly identified neosubstrates in Huh-7 and NB-4 cells according to whether they contained either an IMiD-like or PG core structure, followed by hierarchical clustering of protein fold-changes (i.e., MGD vs. DMSO) within both subsets. Remarkably, this showed that most known IMiD targets such as IKZF1, ZFP91, CSNK1A1, GSPT1, GZF1 or RAB28 were only hit by MGDs with IMiD-like core structures. In contrast, LIMD1 and novel neosubstrates such as KDM4B or FLCN were exclusively targeted by PG-based compounds, which also exhibited higher neosubstrate selectivity (Fig. 4b and Supplementary Fig. 13). A third category of neosubstrates included proteins such as FIZ1, ZBTB39 and the novel neosubstrate G3BP2, which were degraded to a similar extent by both PG-based and IMiD-like MGDs. Remarkably, while the IMiD-like G3BP2 degrader SJ42229 co-depleted several known IMiD targets, none of these were affected by the selective PG-based G3BP2 degrader SJ41824 (Fig. 4c).

**Figure 4.**
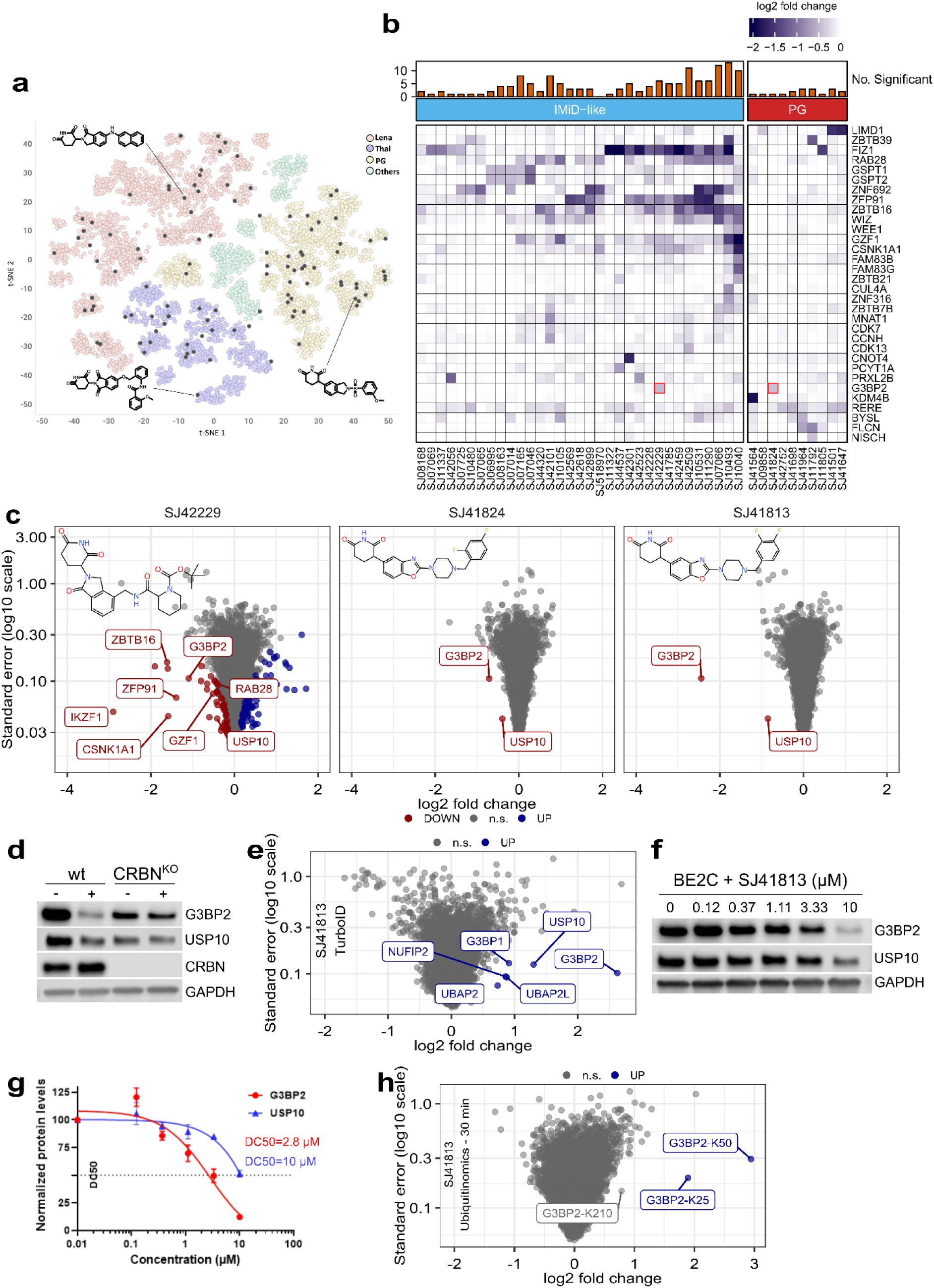
Structure-degradation relationship analysis of screened MGDs. (A) t-SNE plot showing clustering of MGDs according to their core structure. Lena= lenalidomide, Thal= thalidomide, PG= phenyl-glutarimide. Each dot represents a compound, and the black dots denote the analogues selected for proteomics screening (100 compounds). Representative structures for lenalidomide-, pomalidomide- or PG-based cores are shown. (B) Heat map of log_2_ fold changes (MGD vs. DMSO) of known and newly identified neosubstrates in Huh-7. The compounds were arranged according to whether they contained an IMiD-like or a PG core structure. The bar chart on top shows the number of significantly downregulated neosubstrates for each compound. G3BP2 regulation by SJ42229 and SJ41824 is marked by red rectangles. (C) Volcano plots depicting protein quantifications for the G3BP2 degraders SJ42229 (IMiD-like core, left), SJ41824 (PG-based core, middle) and SJ41813 (PG-based core, right). All compounds were tested at a concentration of 10 µM in 24-hour treatments of NB-4 cells. The x-axis shows log2 fold-changes and the standard errors are plotted on the y-axis (log_10_ scale). Significantly up- and downregulated proteins are colored in blue and red, respectively (LIMMA^29^, FDR=1%). IMiD-derived neosubstrates are highlighted for SJ42229. The chemical structures of each degrader are shown within their respective volcano plots. n.s.= not significant. (D) Western blot analysis of the indicated proteins in SK-N-BE(2)-C wild-type and CRBN knockout (CRBN^KO^) cells treated with DMSO or 10 µM of SJ41813 for 24 hours. (E) Volcano plot of the biotinylated proteome as determined by TurboID-MS upon treatment of HEK293 cells expressing TurboID-CRBN with SJ41813. The x-axis shows log_2_ fold-changes and the y-axis standard error on a log_10_ scale. Significantly enriched proteins (LIMMA^29^, FDR=1%) are marked in blue and are labeled. n.s.= not significant. (F) Immunoblots for G3BP2 and USP10 proteins after the treatment of SK-N-BE(2)-C cells with increasing concentrations (in µM) of SJ41813 for 24 hours. Depicted blots are representative of two independent experiments. (G) Quantification of Western blot bands of G3BP2 and USP10 from (F). The half-maximum degradation concentrations (DC_50_) are shown for both proteins. (H) Volcano plot of the ubiquitinome of NB-4 cells treated with SJ41813 for 30 minutes. The x-axis shows log_2_ fold-changes and the y-axis the standard error. Significantly increased ubiquitinated peptides (LIMMA^29^, FDR=5%) are marked in blue and G3BP2 ubiquitination sites are highlighted. n.s.= not significant.

### Comprehensive characterization and chemical optimization of MGDs targeting G3BP2 and KDM4B

Given the advantage of high compound selectivity in defining starting points for further chemical optimization, we selected the PG-based MGD SJ41824 targeting G3BP2 for chemical analog testing. G3BP2 is an mRNA binding protein that physically associates with both G3BP1 and USP10, playing a key role in protecting ribosomes from degradation under cellular stress^48^. At present, no selective G3BP2 inhibitors have been identified. To establish structure-degradation relationship around this chemotype and potentially identify more potent G3BP2 degraders, we evaluated additional analogs of the G3BP2 degrader SJ41824 in NB-4 cells by global proteomic profiling. Among these, SJ41813 caused stronger and highly selective degradation of both G3BP2 and USP10, as determined by proteomics and Western blotting (Fig. 4c and Supplementary Fig. 14a,b). Intriguingly, SJ41813 differs from its regioisomer SJ41824 solely by the presence of a 3,4-difluorophenyl instead of a 2,4-difluorophenyl group, highlighting that even minor structural modifications can have a strong impact on degradation efficacy and potency^49^. CRISPR-Cas9-mediated knockout of CRBN in SK-N-BE(2)-C cells completely abrogated G3BP2 degradation by SJ41813 (Fig. 4d), confirming a CRBN-dependent mechanism. To demonstrate ternary complex formation in cells between SJ41813-engaged CRBN and G3BP2, we engineered a HEK293 cell line with doxycycline-inducible expression of TurboID-CRBN^50^. We treated these cells with the degrader, followed by affinity-purification of biotinylated proteins and quantitative mass spectrometry. Strikingly, this revealed a >6-fold enrichment of G3BP2, along with a weaker enrichment of its binding partners, including USP10 (Fig. 4e). This clearly pointed to G3BP2 as direct CRBN neosubstrate, which was further corroborated by a higher degradation potency of G3BP2 compared to USP10, with DC_50_ values of 2.8 µM and 10 µM, respectively (Fig. 4f,g). Moreover, quantitative ubiquitinomics revealed two upregulated ubiquitination sites on G3BP2 in SJ41813-treated NB-4 cells but none on USP10 (Figure 4h). This was further corroborated by a significantly higher ubiquitination of G3BP2 compared to USP10 in SJ41813-treated HEK293-CRBNoe cells (Supplementary Fig. 14). Collectively, these data clearly point to G3BP2 as the direct target, while USP10 acts as bystander neosubstrate.

Another PG-based degrader selected for chemical analog testing was SJ41564, which strongly and selectively downregulated the protein levels of KDM4B. KDM4B is a histone demethylase that specifically acts on lysine 9 and lysine 36 of histone H3 (H3K9 and H3K36), which are playing a key role in regulation of gene expression and chromatin organization. It shares high homology with the five KDM4 family members, including KDM4A and KDM4C, that are frequently overexpressed in cancer^51^. Despite considerable efforts across the pharmaceutical industry and academia^52^, selective inhibitors for individual KDM4 family members have not been identified, mostly due to the high structural similarity of their catalytic JmjC domains. Intriguingly, SJ41564 selectively degraded KDM4B without affecting KDM4A, KDM4C or KDM4D levels in NB-4 and Huh-7 (Fig. 5a). The expression of KDM4A was slightly increased following KDM4B degradation in SK-N-BE(2)-C cells, indicating a potential compensatory effect specific to this neuroblastoma cell line (Supplementary Fig. 15a). Indeed, a similar compensatory KDM4A upregulation has been shown responsible to mitigate the effects of genetic silencing of KDM4B on the growth of rhabdosarcoma cells^53^. To further validate the mechanism of action of SJ41564 as a CRBN-dependent MGD, we performed a side-by-side comparison with SJ48108, an N-methylated negative control. As expected, treatment of SK-N-BE(2)-C cells with SJ41564, but not SJ48108, led to a rapid and potent KDM4B degradation, with a DC_50_ of 36 nM and a D_max_ of 92% at 10 µM. (Fig. 5b-c and Supplementary Fig. 15b). In addition to HEK293, Huh-7, NB-4 and SK-N-BE(2)-C, we observed KDM4B degradation in six other cancer cell lines (Supplementary Fig. 15c). Consistent with proteomics analyses in NB-4 cells, SJ41564-mediated KDM4B degradation in SK-N-BE(2)-C cells was completely prevented by co-treatment with either the proteasomal inhibitor MG132, the NEDD8 inhibitor MLN4924, or the CRBN ligand lenalidomide. Furthermore, degradation of KDM4B by SJ41564 was abolished in SK-N-BE(2)-C CRBN^KO^ cells and was not affecting its mRNA levels. This demonstrates a mechanism of action that is dependent on CRBN and on the ubiquitin-proteasome pathway (Supplementary Fig. 15d-g).

**Figure 5.**
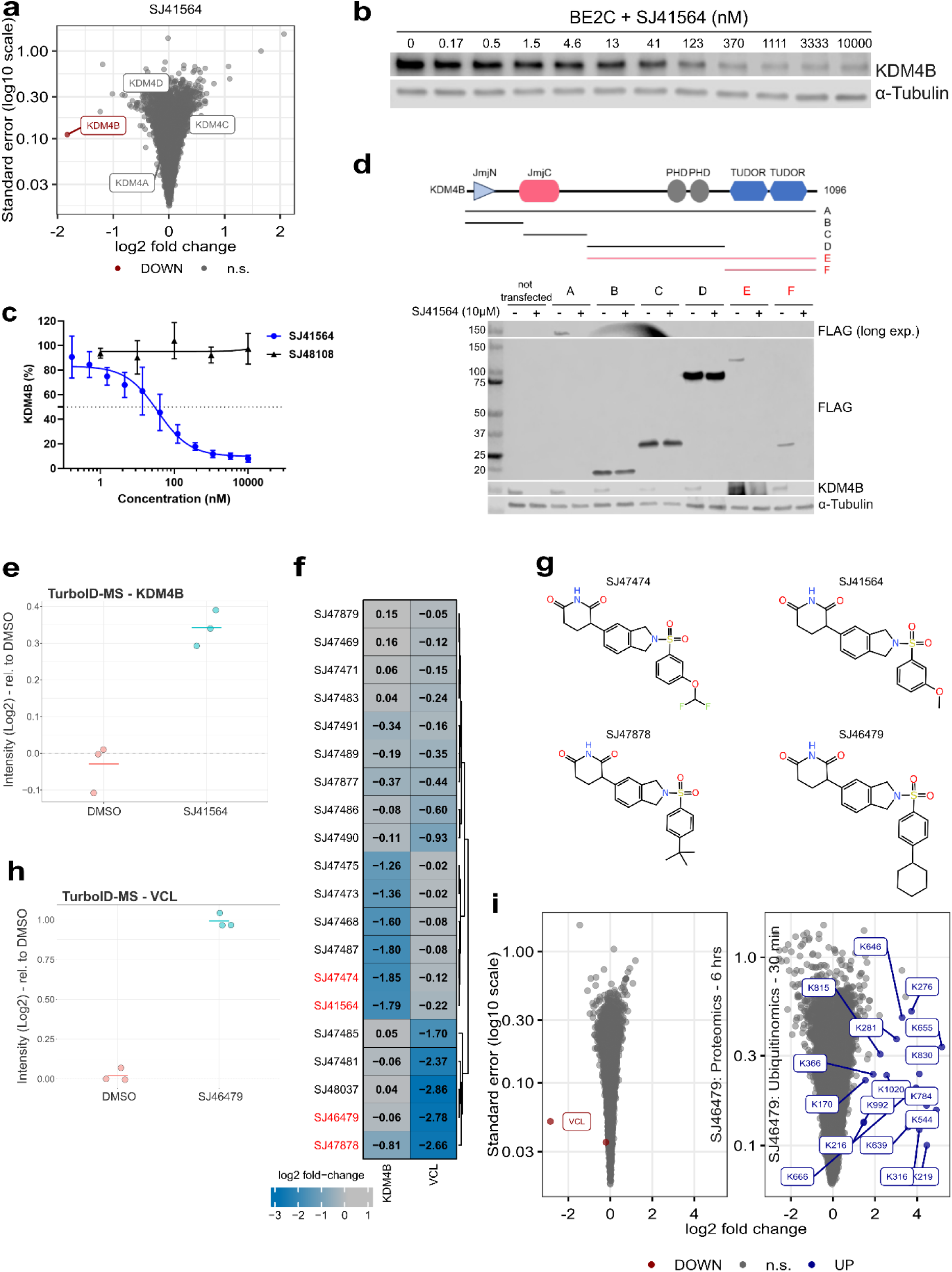
In-depth characterization of KDM4B and VCL as CRBN neosubstrates. (A) Volcano plot visualization of the proteome for SJ41564-treated NB-4 cells (6 hours). The x-axis shows log_2_ fold-changes and the y-axis the standard error on a log_10_ scale. Quantified KDM4 family members are highlighted, with KDM4B being significantly downregulated (in red). n.s.= not significant. (B) Immunoblots for KDM4B and α-tubulin proteins after treatment of SK-N-BE(2)-C cells with increasing concentrations (in nM) of SJ41564 for 24 hours. Depicted blots are representative of three independent experiments. (C) Quantification of KDM4B Western blot for SJ41564 shown in (B). The N-methylated negative control compound SJ48108 was included in the analysis. (D) Top: Protein architecture of KDM4B and schematic of truncated KDM4B protein fragments. Bottom: SK-N-BE(2)-C cells overexpressing CRBN were transiently transfected with Flag-tagged KDM4B fragments A-F. Depicted are immunoblots for FLAG-tagged KDM4B fragment proteins after treating cells with 10 µM of SJ41564 for 24 hours. (E) KDM4B levels as quantified by TurboID-MS of HEK293-TurboID-CRBN cells treated with bortezomib (0.5 µM) or the KDM4B degrader SJ41564 along with bortezomib for 120 minutes. KDM4B was significantly enriched (LIMMA^29^, FDR=1%). (F) Heat map of VCL and KDM4B protein log_2_ fold-changes (degrader vs. DMSO) quantified by global proteomics for various MGD compounds. (G) Chemical structures of selected KDM4B and VCL degrader compounds. (H) VCL levels as quantified by TurboID-MS of HEK293-TurboID-CRBN cells treated with bortezomib (0.5 µM) or the VCL degrader SJ41564 along with bortezomib for 120 minutes. VCL was significantly enriched (LIMMA^29^, FDR=1%). (I) Volcano plot visualization of the proteome (6-hour treatment) and the ubiquitinome (30-minute treatment) for SJ46479-treated NB-4 cells. The x-axis shows log_2_ fold-changes and the y-axis the standard error on a log_10_ scale. Significantly up- or downregulatedproteins ((LIMMA^29^, FDR=1%) or ubiquitinated peptides (LIMMA^29^, FDR=5%) are marked in blue and red, respectively. Ubiquitination sites mapping to VCL are highlighted. n.s.= not significant.

After establishing the CRBN dependency, we endeavored to map the region of KDM4B that interacts with CRBN-bound SJ41564. To do that, we generated a series of KDM4B fragments (A-F) and assessed their degradation by Western blotting in SK-N-BE(2)-C cells overexpressing CRBN following treatment with the MGD. Treatment of SJ41564 selectively induced potent degradation of fragments E and F, clearly pinpointing the Tudor domains at the less conserved C-terminus as responsible for mediating KDM4B degradation (Fig. 5d).

TurboID-MS experiments confirmed the SJ41564-induced KDM4B-CRBN complex formation in intact HEK293 cells (Fig. 5e). Subsequent evaluation of SJ41564 analogs by quantitative proteomics revealed several additional potent KDM4B degraders and identified substitution at the *meta-*position of the distal phenyl moiety as preferred for cellular activity (Fig. 5f, Supplementary Fig.16).

Strikingly, while analogs substituted at the *para-*position showed little or no KDM4B degrading activity, we were intrigued to find that several of these PG-based CRBN binders caused dramatic reductions in the key cytoskeletal protein VCL^54^. For example, SJ46479 and SJ47878 reduced VCL levels by 90% within 6 hours with high selectivity in NB-4 and Caki-1 cells (Fig. 5f,g). We further established time-, dose- and CRBN-dependent VCL degradation by SJ46479 in Caki-1 cells where it displayed a DC_50_ value of 255 nM and a D_max_ of 96% (Supplementary Fig. 17). Finally, using biotin proximity labeling-MS, we demonstrated intracellular MGD-induced complex formation with CRBN, and we detected rapid ubiquitination of over 20 lysine residues on VCL following treatment of NB-4 cells with SJ46479 (Fig. 5h,i). In summary, our proteomics screening approach identified a number of exquisitely selective PG-based degraders for G3BP2, KDM4B and VCL. Our work further exemplifies how subtle chemical variations in CRBN-based degraders can affect MGD neosubstrate specificity, resulting in highly efficacious and selective degraders for previously unknown targets.

## Conclusions

In this study, we report a highly sensitive and unbiased proteomics screening workflow for discovery of MGDs and their targets in intact cells. Our platform employs the testing of compounds across multiple time points and cell lines, utilizes neddylation inhibition to confirm direct and CRBN-dependent protein degradation events, and includes rigorous target validation through ubiquitinomics. Screening of a subset of 100 diverse CRBN ligands enabled us to discover and validate 50 previously unreported neosubstrates. We also validated numerous neosubstrates for which only ternary complex formation has been previously observed in biochemical or cell-based assays^9,35,36,40^. To enhance the sensitivity of our screening platform we utilized HEK293 cells overexpressing CRBN, which enabled us to confirm weakly degraded putative neosubstrates such as ZBTB21 and CSNK1E, initially identified in the primary Huh-7 and NB-4 screens.

Importantly, our integrated proteomics screening workflow rapidly generates high-quality data, facilitating not only hit identification and validation but also supporting subsequent chemical optimization efforts^40,55^. For example, phenyl glutarimide SJ41824 was identified as a potent degrader of G3BP2, a stress granule-associated protein with a potential role in the development of breast and prostate cancers^56,57^. Given that no inhibitors for G3BP2 have been reported, we synthesized several analogues of SJ41824 and evaluated them using our proteomics platform, which allowed us to further validate our initial findings and establish preliminary structure-degradation relationships. This led to the discovery of SJ41813, a closely related regioisomeric analogue of SJ41824 displaying more potent degradation of G3BP2, with a DC_50_ value of 2.8 µM. Notably, the quantitative ubiquitinomics component of our screening workflow demonstrated that USP10, the only other significantly downregulated protein by SJ41813, was a bystander target.

Interestingly, another phenyl glutarimide analogue from our screening set, SJ41564, was found to be a potent and selective degrader of KDM4B, a histone demethylase that for many years attracted a lot of interest in cancer drug discovery. However, despite multiple and extensive efforts, developing selective KDM4B inhibitors proved to be highly challenging, mostly because they were designed to target the highly conserved enzyme pocket and compete with a highly abundant co-factor^58^. And yet, while inducing a highly potent CRBN-dependent degradation of KDM4B with DC_50_ of 36 nM and 92% Dmax, SJ41564 did not significantly affect any of the over 9,000 quantified proteins in NB-4 cells after a 6-hour incubation at a concentration of 10 µM. Using a range of Flag-tagged constructs we were able to identify the less conserved Tudor domains as the mediators of the KDM4B degradation, which could potentially rationalize the impressive selectivity of SJ41564. Remarkably, as evaluation of SJ41564 analogs by quantitative proteomics revealed several additional potent KDM4B degraders, it also led to the discovery of potent and highly selective degraders of the cytoskeletal protein VCL, such as SJ46479 (VCL DC_50_= 255 nM; D_max_ = 96%). These findings align well with our earlier observations of rather steep structure-degradation relationships that appear to be characteristic for molecular glue degraders.

In summary, our unbiased high-throughput proteomics screening approach identified a number of exquisitely potent, selective and structurally novel phenyl glutarimide-based degraders of G3BP2, KDM4B and VCL proteins. Interestingly, none of these novel neosubstrates contain the classical degron motif, suggesting that the proteome accessible to CRBN-dependent modulation is greater and more diverse than the one defined by the original IMiDs. While the underlying structural determinants remain to be identified, this study provides a strong rationale for proteomic screening of diverse compound libraries built around novel scaffolds, to discover neosubstrates in the disease-relevant, formerly undruggable target space.

## Supplementary Figures

**Figure S1:**
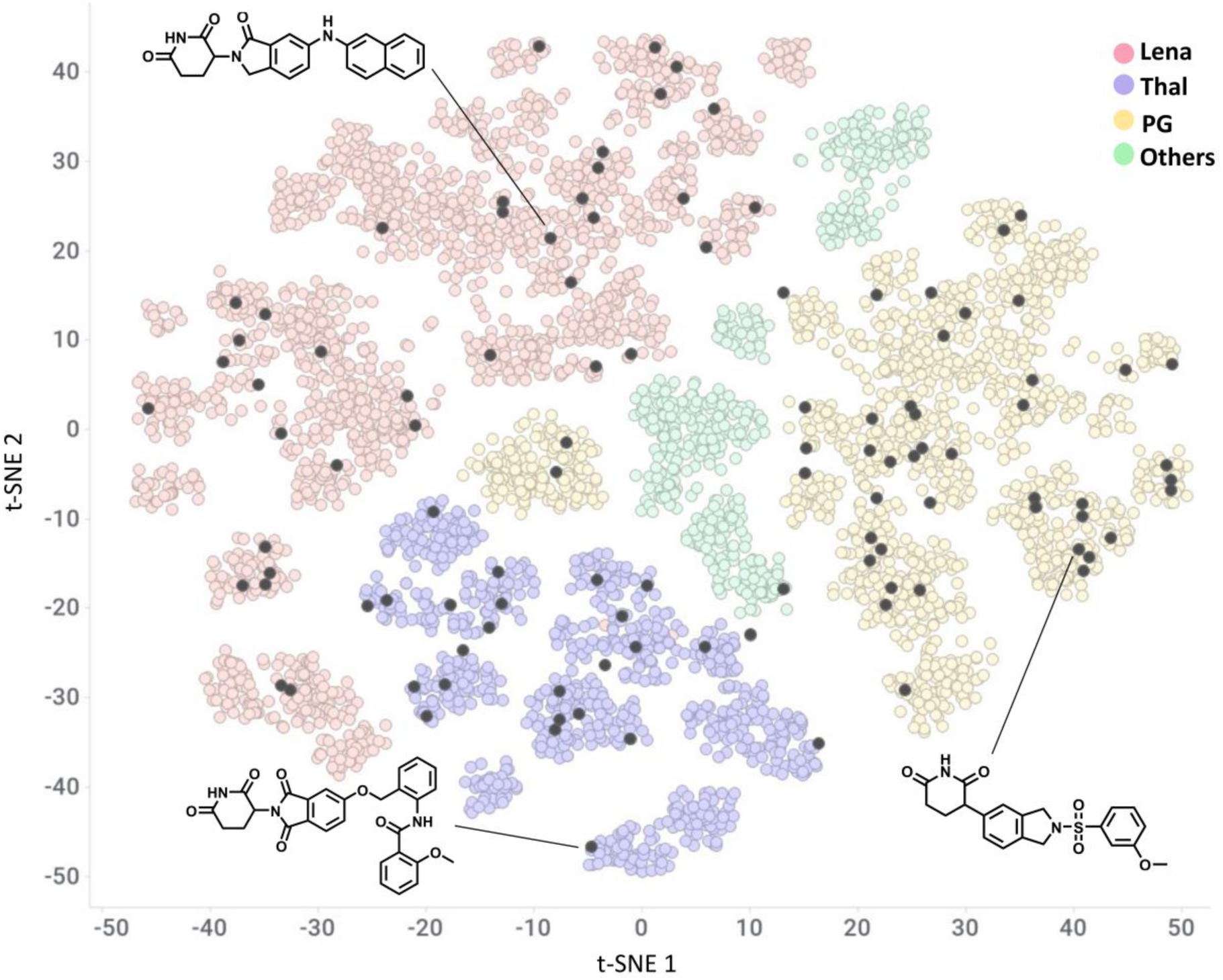
t-SNE plot representation of the molecular glue library comprising 5,000 compounds. The clustering is based on molecular fingerprints and the colors are coded based on the CRBN-binding core diversity (lenalidomide, thalidomide, phenyl-glutarimide (PG), others). Each dot represents a compound, and the black dots denote the analogues selected for proteomics screening (100 compounds). Representative structures for lenalidomide-, pomalidomide- or PG-based cores are shown.

**Figure S2.**
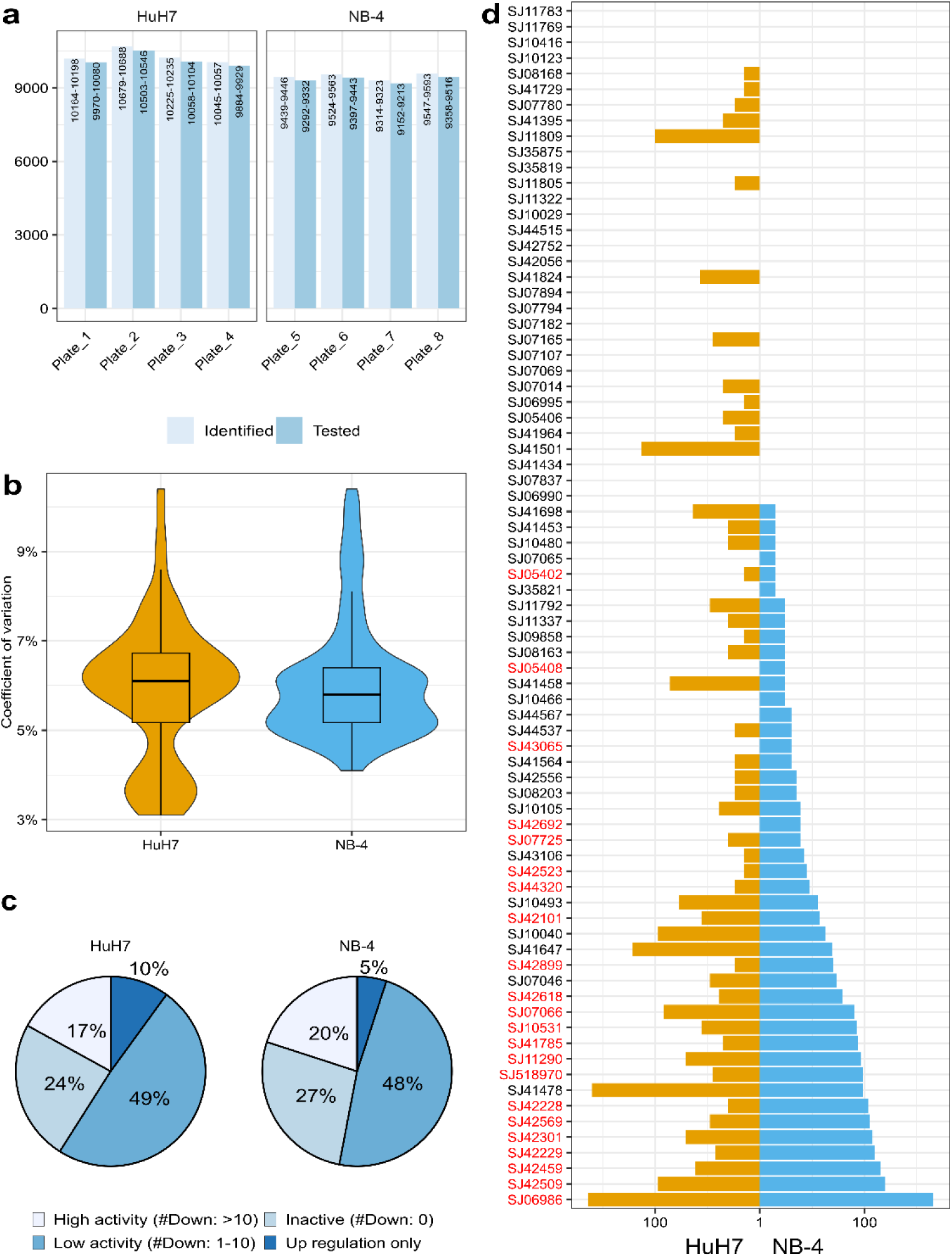
Summary of proteomics screening data from Huh-7 and NB-4 cells. (A) Bar chart displaying numbers of all proteins quantified in Huh-7 and NB-4 cells (24-hour compound screening) after filtering for missing values (light blue) and the subset of proteins that underwent statistical testing (darker blue), which required 100% data completeness in the compound-treated samples. (B) Violin plot showing distribution of protein coefficients of variation (%) in both cell lines. © Pie chart displaying the activity of all screened compounds, defined by the number of induced significant (LIMMA^29^, FDR=1%) protein regulations. Inactive = no significantly modulated protein; Low activity = number of regulated proteins <10; High activity = number of regulated proteins >10; In dark blue are compounds that did not induce any protein downregulations but significant protein upregulations. (D) Number of significantly downregulated proteins in Huh-7 and NB-4 upon screening of 100 MGD compounds following 24 hours of treatment. Degraders of IKZF1 are marked in red.

**Figure S3.**
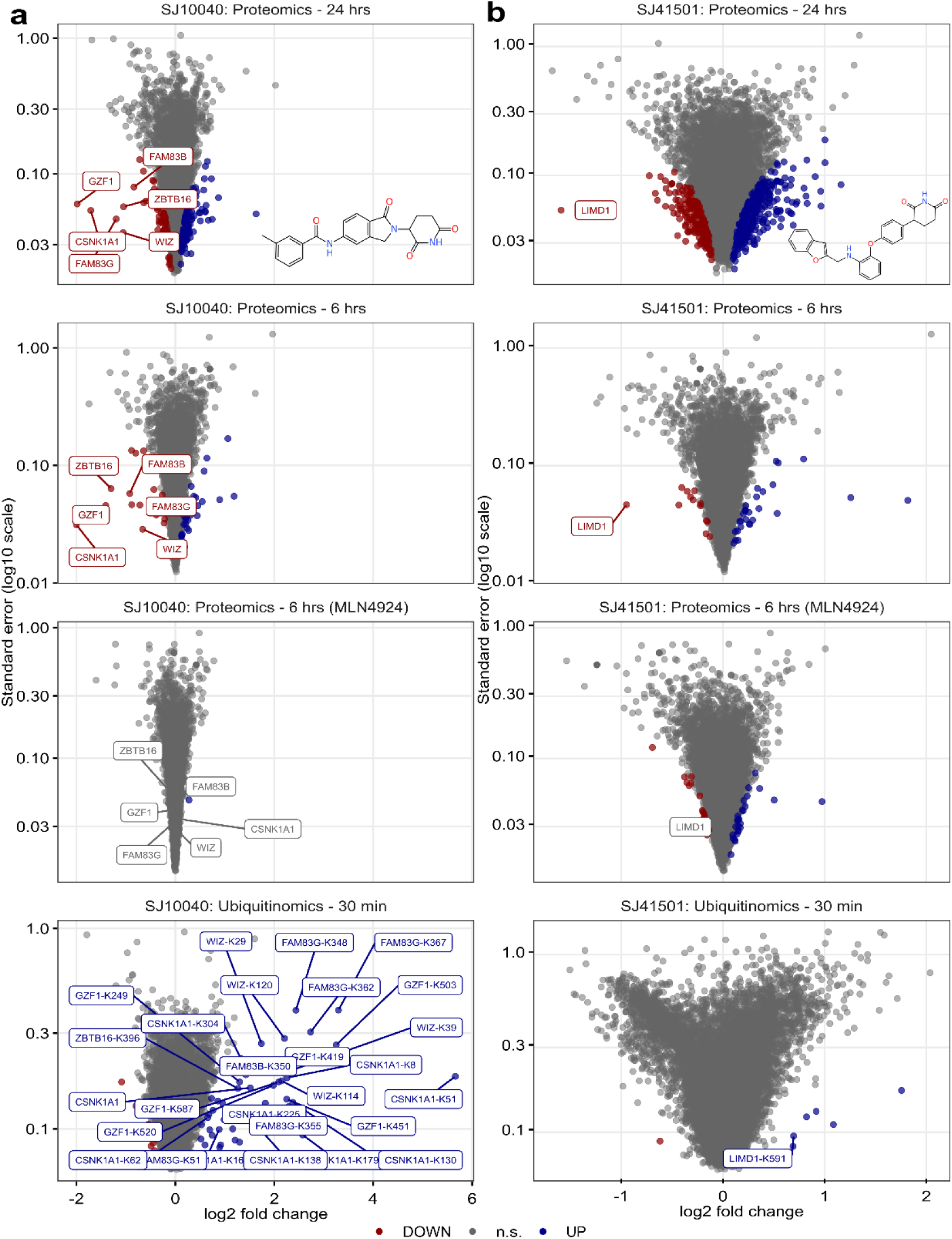
Examples of previously reported neosubstrates validated in our screen. (C) and (B) Volcano plots of the proteomes after treatment with the indicated compounds for 24 hours (top), 6 hours (second from top), 6 hours in the presence of MLN4924 (second from bottom), and the ubiquitinome following 30 minutes of compound treatment (bottom). The x-axis shows log-transformed fold-changes and the y-axis the standard error on a log10 scale. Previously reported neosubstrates are highlighted, with significantly upregulated proteins (LIMMA^29^, FDR=1%) marked in blue and downregulated proteins marked in red. n.s.= not significant.

**Figure S4.**
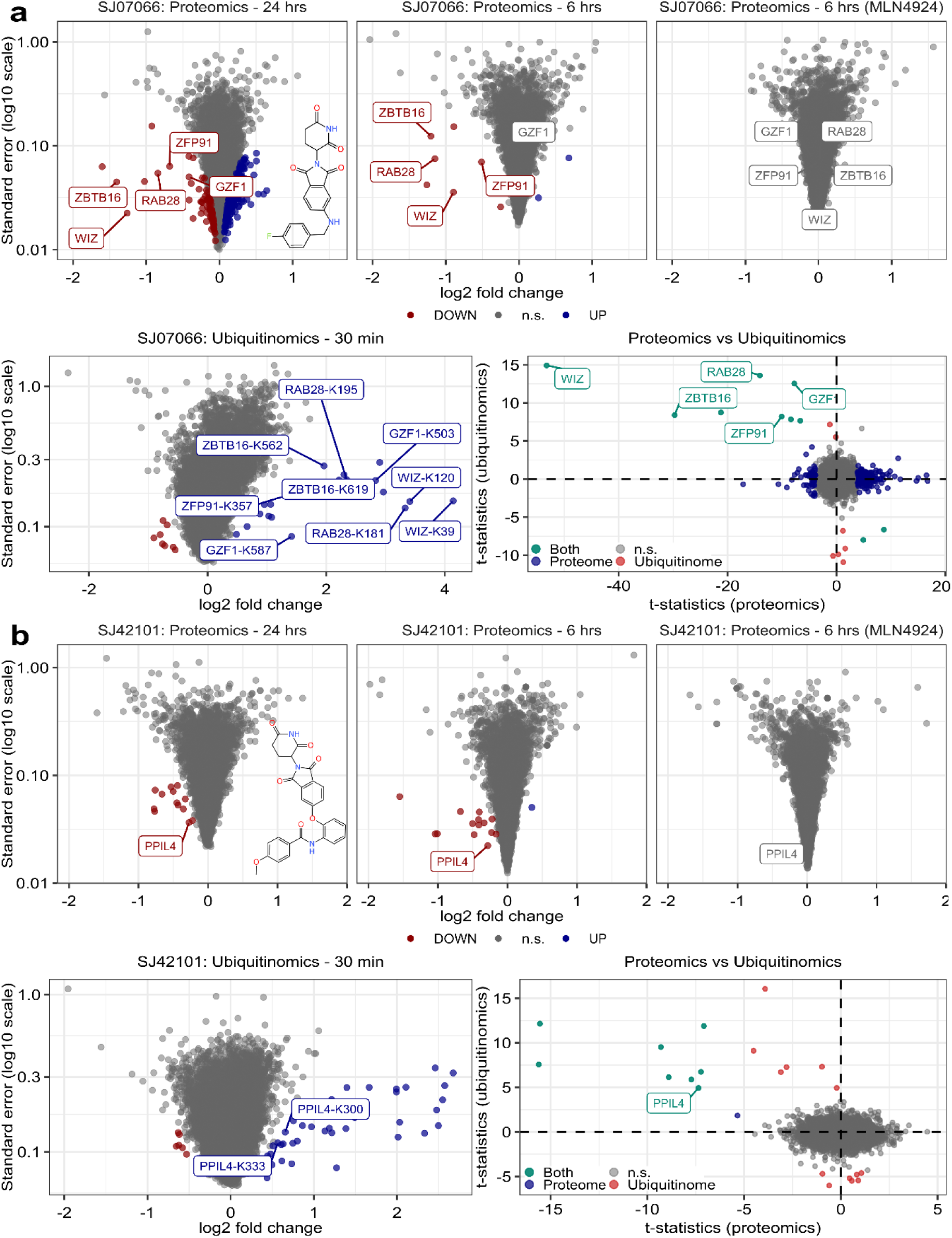
Examples of previously reported neosubstrates validated in our screen. (C) and (B) Volcano plots of the proteomes are presented following treatment with the indicated compounds for 24 hours (top left), 6 hours (top middle), and 6 hours in the presence of MLN4924 (top right). The ubiquitinome following 30 minutes of compound treatment is shown in the bottom left, while the bottom right displays a scatter plot comparing the proteomics and ubiquitinomics t-statistics. Previously reported neosubstrates are highlighted, and significantly upregulated and downregulated proteins (LIMMA^29^, FDR = 1%) are marked in blue and red, respectively. The t-statistics of significantly upregulated ubiquitination sites mapping to the same proteins were averaged. In the t-statistic comparison plots, significantly modulated proteins are labeled in blue, ubiquitinated proteins in red, and proteins that are both significantly downregulated following a 24-hour compound treatment and harbor upregulated ubiquitination sites (30-minute compound treatment) are indicated in turquoise. n.s. = not significant.

**Figure S5.**
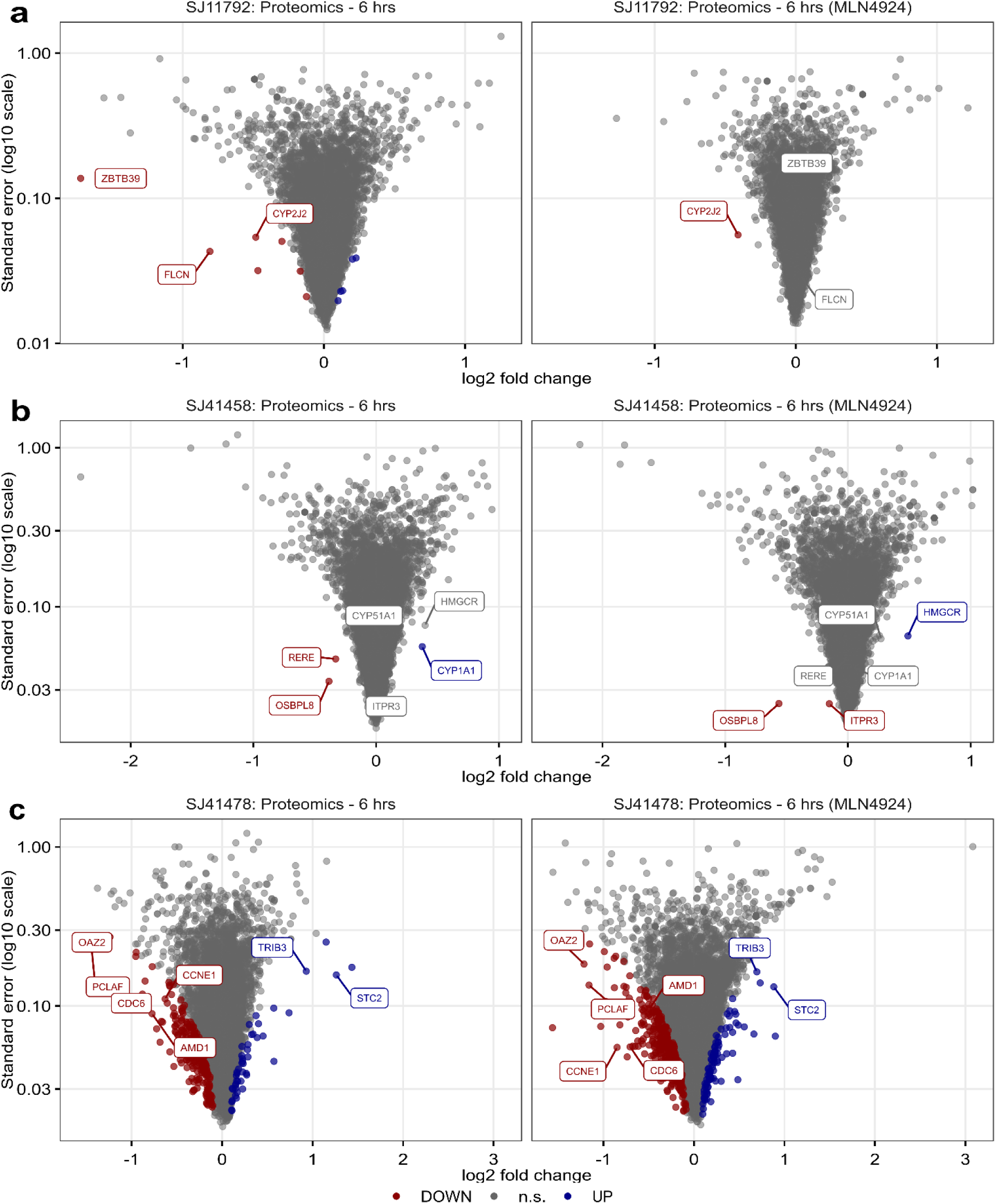
MLN4924-independent protein regulations. (C) – (C) Volcano plots of the proteomes following treatment with the indicated compound for 6 hours, in the absence (left) or presence (right) of MLN4924. Significantly upregulated and downregulated proteins (LIMMA^29^, FDR = 1%) are marked in blue and red, respectively. Selected proteins are highlighted to illustrate both neddylation-dependent and neddylation-independent regulatory events. n.s. = not significant.

**Figure S6.**
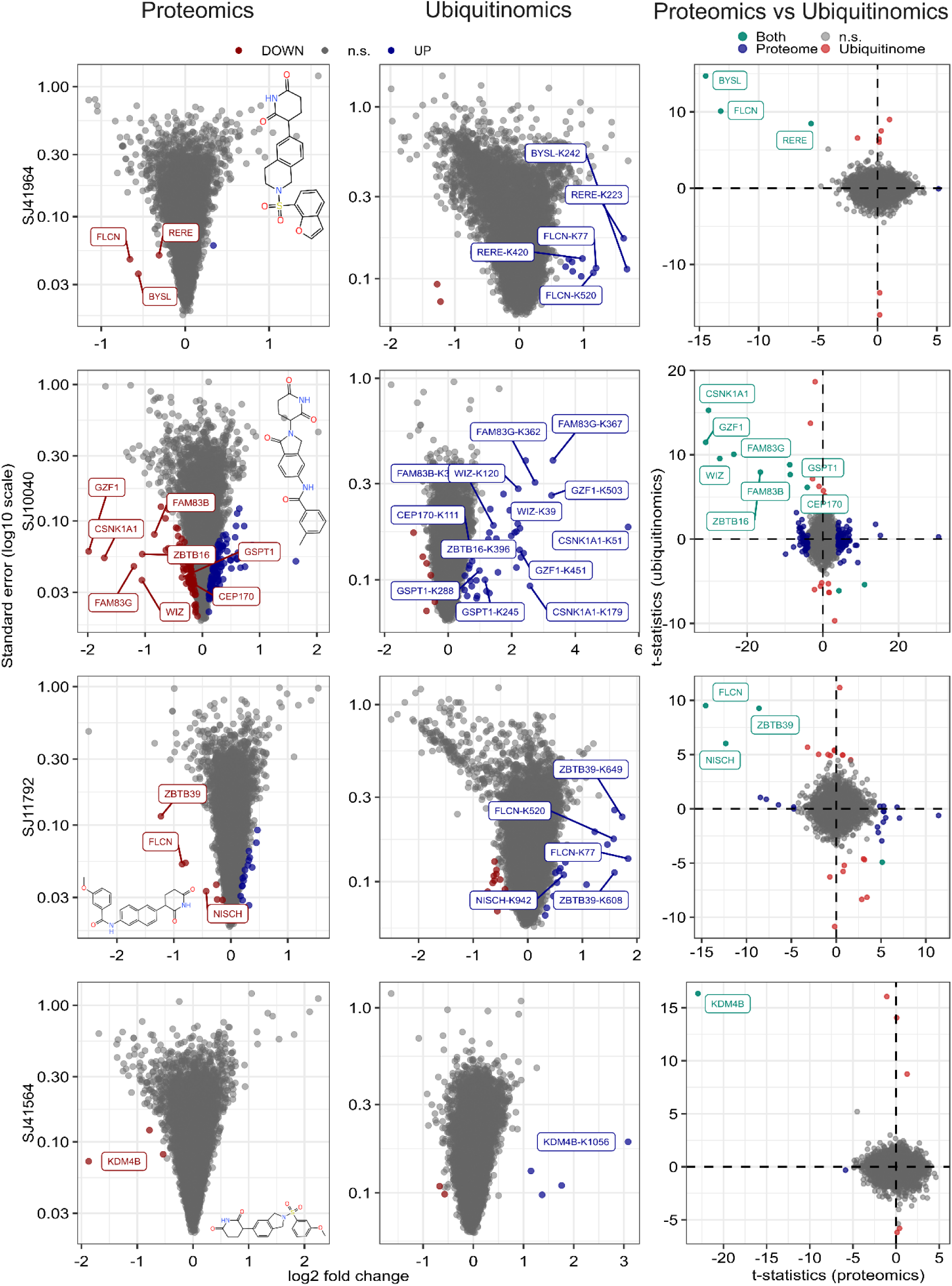
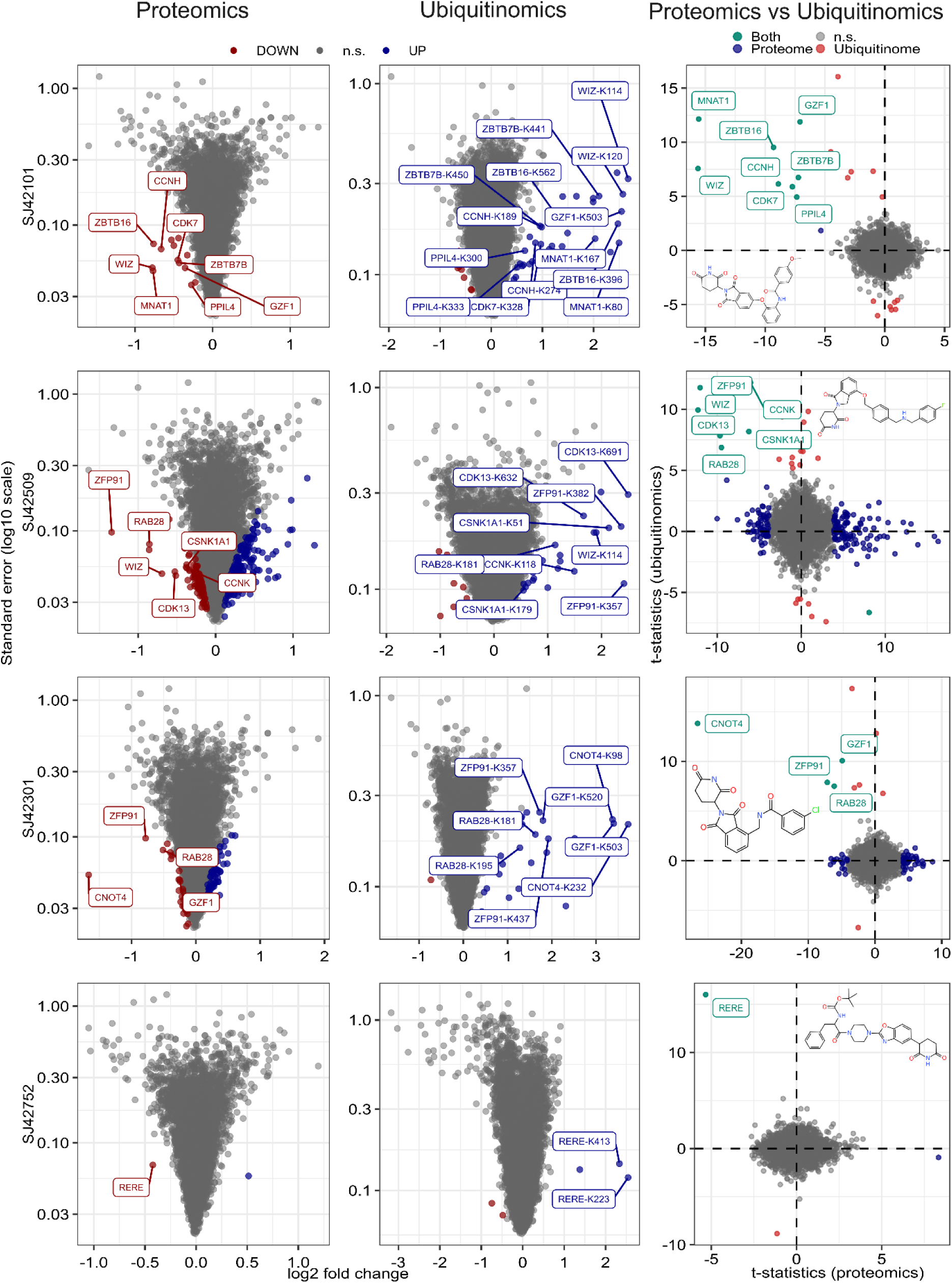
Proteomics and ubiqutinomics screening results leading to the identification of 16 novel neosubstrates in Huh-7. The 24-hour proteomics screening results for each of the indicated compounds are presented on the left. The middle section displays the corresponding ubiquitinomics volcano plots (30 minute treatments), while the right section features a comparison plot of t-statistics between proteomics and ubiquitinomics. In the volcano plots, significantly upregulated proteins or ubiquitinated peptides are indicated in blue, and significantly downregulated proteins or ubiquitinated peptides are indicated in red (LIMMA^29^, FDR = 1% and FDR=5%, for proteomes and ubiquitinomes, respectively). T-statistics for significantly upregulated ubiquitination sites that map to the same proteins were averaged in the comparison plots. In these plots, significantly modulated proteins are labeled in blue, ubiquitinated proteins are labeled in red, and proteins that are both significantly downregulated and exhibit upregulated ubiquitination sites (or vice versa) are labeled in turquoise. n.s. = not significant.

**Figure S7.**
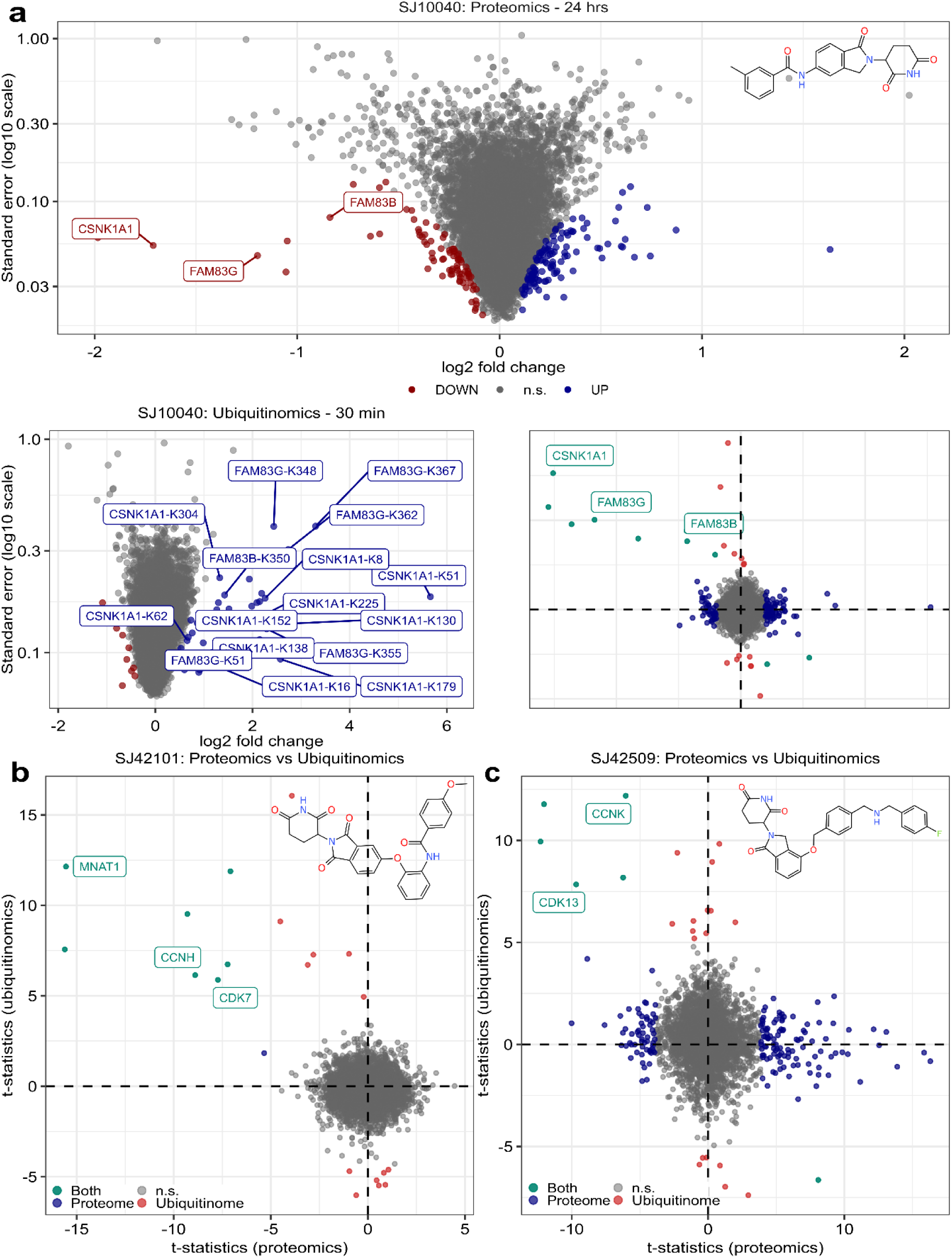
Examples of neosubstrate protein complexes. (A) Volcano plot (top) of the proteome for SJ10040 following treatment for 24 hours, with CSNK1A1, FAM83G and FAM83B significantly downregulated (among others). For each protein, the log_2_-transformed fold-change is depicted on the x-axis and its standard error (log_10_ scale) on the y-axis. Significantly up- and downregulated proteins are marked in blue and red, respectively (LIMMA^29^, FDR = 1%). Volcano plot (bottom left) of the ubiquitinome for SJ10040 (30 minutes of treatment), with CSNK1A1, FAM83G and FAM83B ubiquitination sites significantly upregulated. Significantly up- and downregulated ubiquitinated peptides are marked in blue and red, respectively (LIMMA^29^, FDR = 5%). On the bottom right, a proteomics vs. ubiquitinomics t-statistic comparison plot is shown. Significantly modulated proteins are labeled in blue, ubiquitinated proteins in red and proteins that are both significantly downregulated and harbor upregulated ubiquitination sites are in turquoise. n.s. = not significant. (B) and (C) Proteomics vs. ubiquitinomics t-statistic comparison plots for SJ42101 (B) and SJ42509 (C). T-statistics of significantly upregulated ubiquitination site mapping to the same proteins were averaged. Significantly modulated proteins (LIMMA^29^, FDR = 1%, 24-hour treatment) are labeled in blue, ubiquitinated proteins (LIMMA^29^, FDR = 5%, 30-minute treatment) in red and proteins that are both significantly downregulated and harbor upregulated ubiquitination sites (or vice versa) are in turquoise.

**Figure S8.**
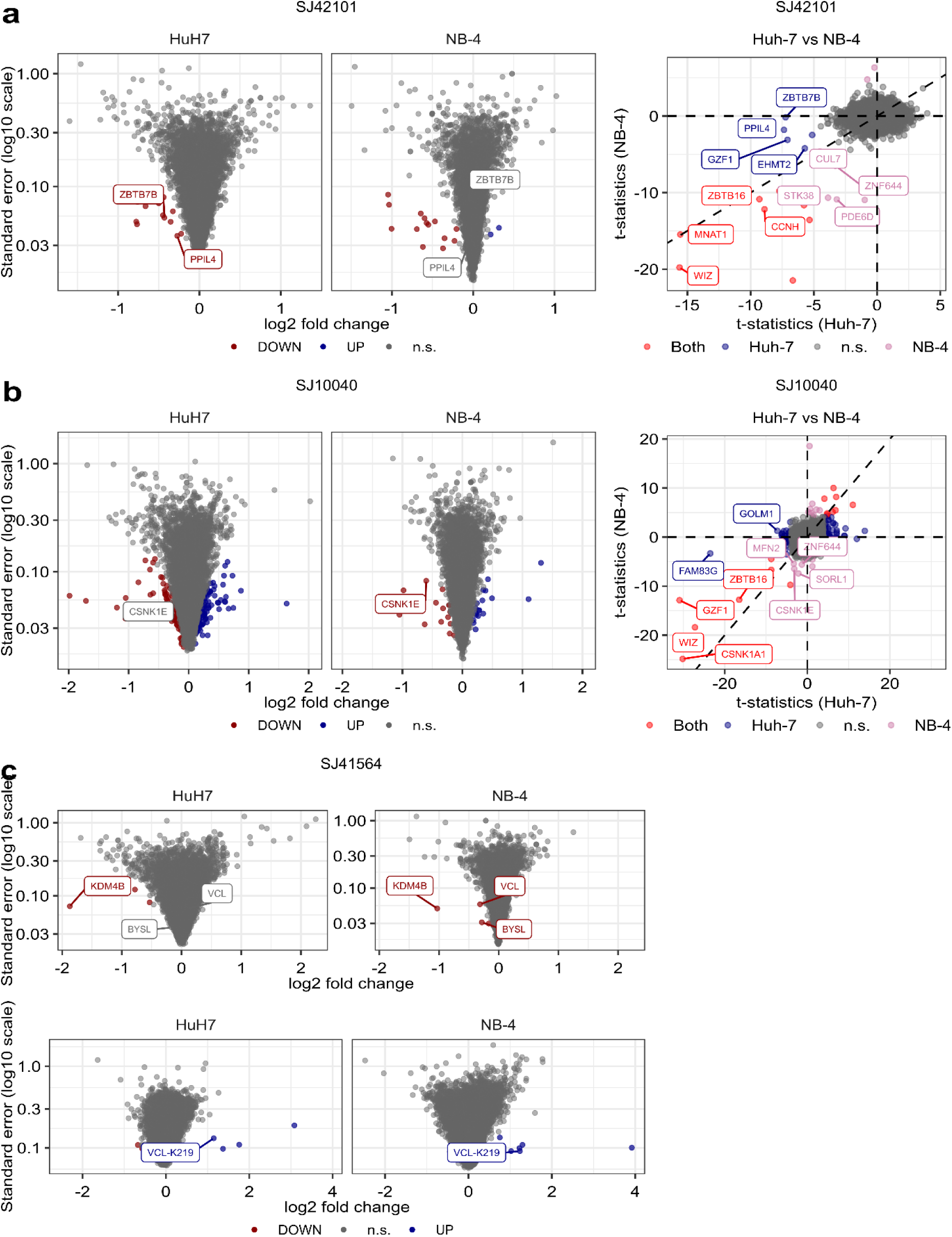
Comparison of proteomic profiles of Huh-7 and NB-4 cells. (A) and (B) Volcano plots for the indicated 24-hour compound treatments in Huh-7 (left) and NB-4 cells (middle). For each protein, the log_2_-transformed fold-change is depicted on the x-axis and its standard error (log_10_ scale) on the y-axis. Significantly up- and downregulated proteins (LIMMA^29^, FDR = 1%) are marked in blue and red, respectively. A t-statistic comparison plot is shown on the right. Significantly modulated proteins in NB-4 cells are labeled in purple, significantly modulated proteins in Huh-7 cells in blue and proteins that are significantly modulated in both cell lines are marked in red. n.s. = not significant. (C) Volcano plots of SJ41564-treated Huh-7 (left) and NB-4 (right). The treatment was for 24 hours. A volcano plot depicting ubiquitinated peptides for both cell lines is shown below. One VCL site was significantly induced and is marked in blue. Significantly up- and downregulated ubiquitinated peptides (LIMMA^29^, FDR = 5%) are marked in blue and red, respectively. n.s. = not significant.

**Figure S9.**
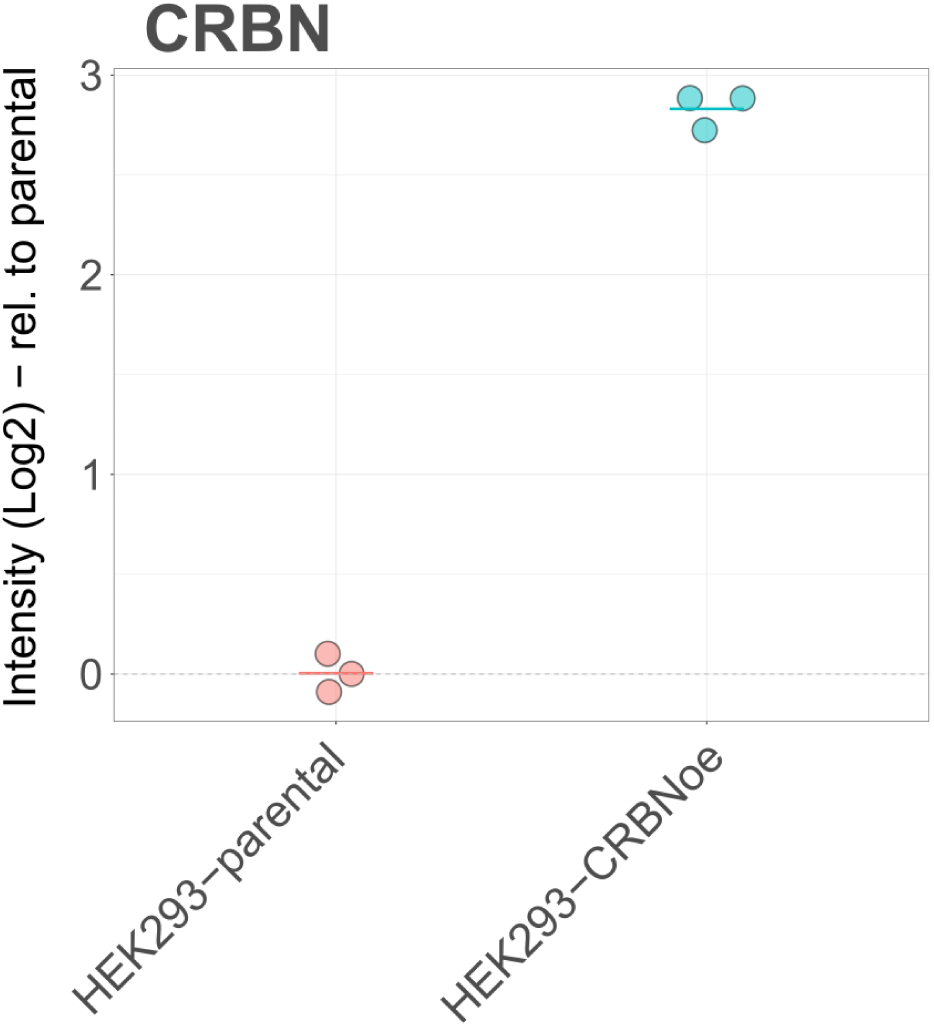
MS-based quantification of CRBN levels in HEK293-CRBNoe cells. Scatter plot illustrating log_2_-transformed CRBN levels quantified by mass spectrometry (MS)-based proteomics in HEK293 cells and HEK293 cells overexpressing CRBN (HEK293-CRBNoe).

**Figure S10.**
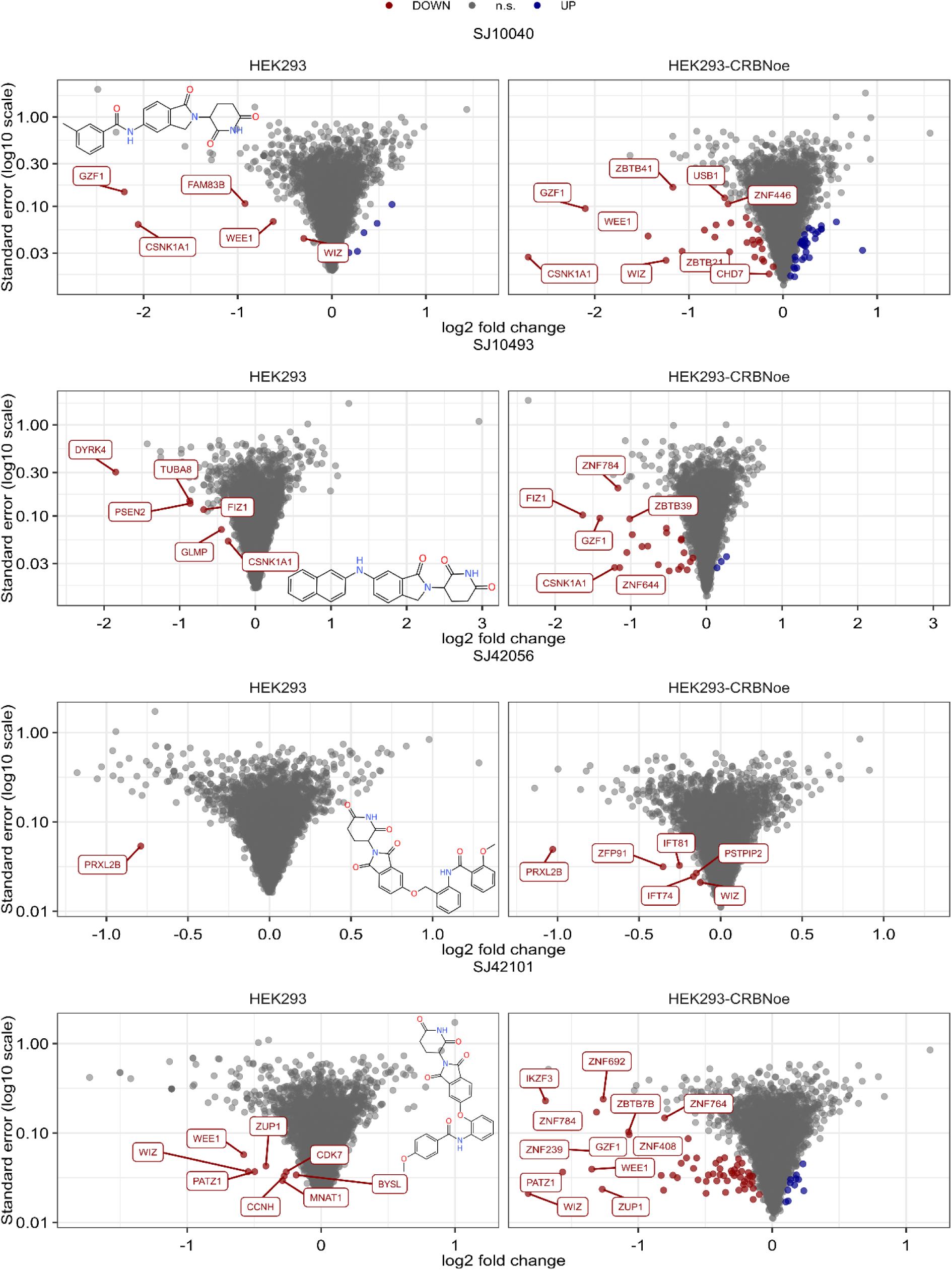
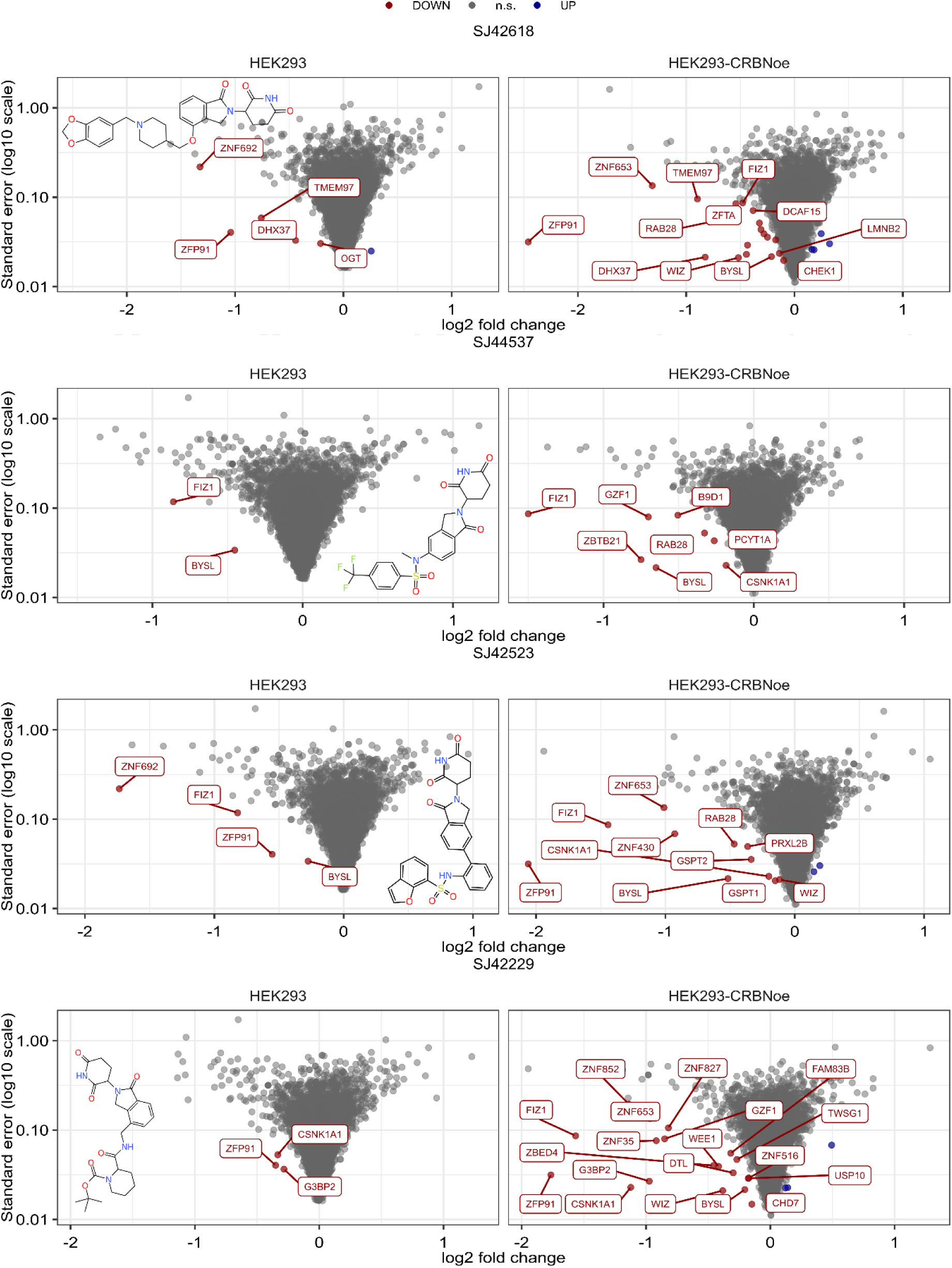
Characterization of HEK293-CRBNoe cells for the responsiveness to MGDs. Volcano plot for the indicated 6-hour compound treatments in HEK293 (left) and HEK293-CRBNoe cells (right). The log_2_-transformed fold-change is depicted on the x-axis and the standard error (log_10_ scale) on the y-axis. Significantly up- and downregulated proteins (LIMMA, FDR = 1%) are marked in blue and red, respectively. n.s. = not significant.

**Figure S11.**
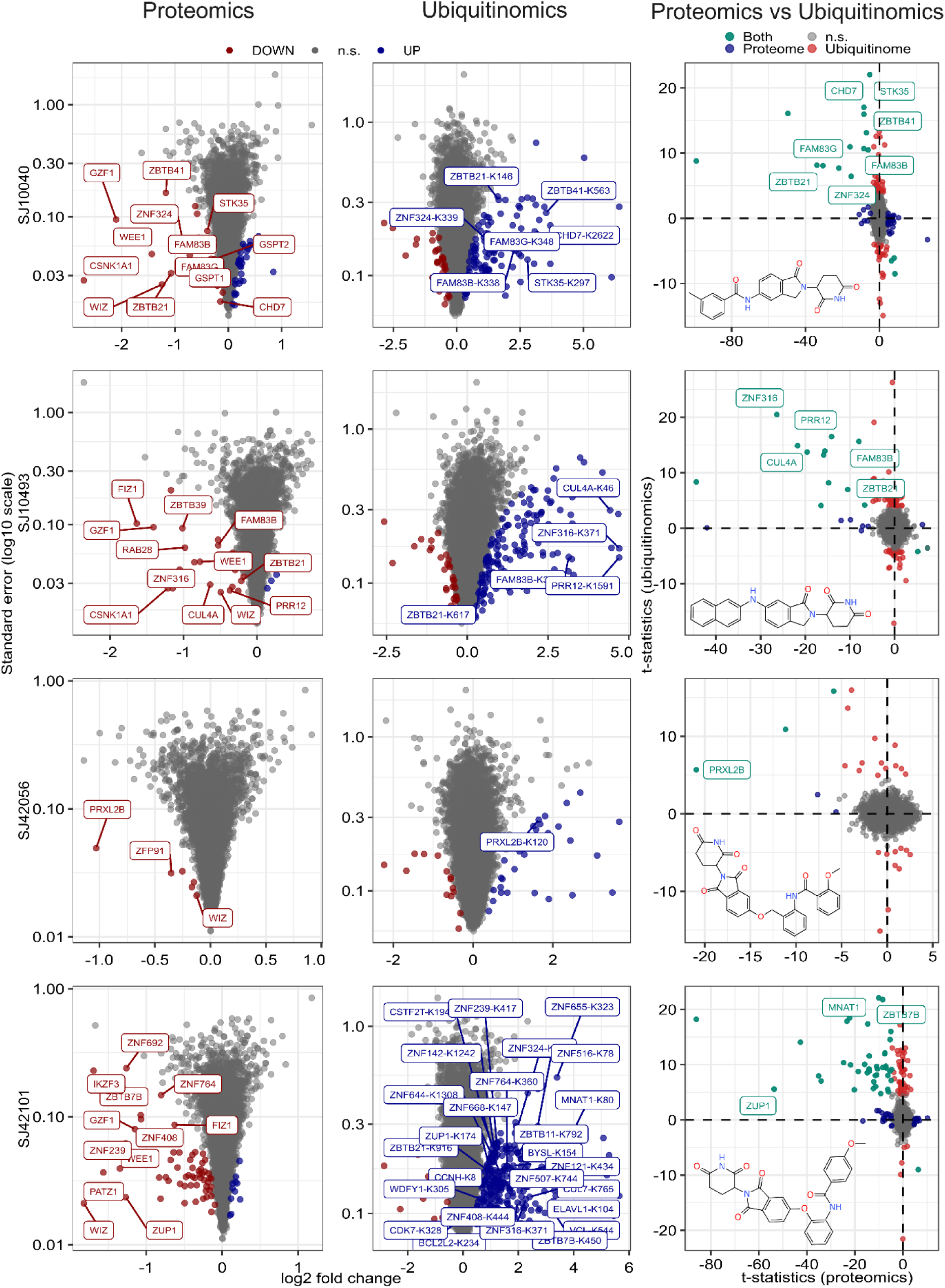
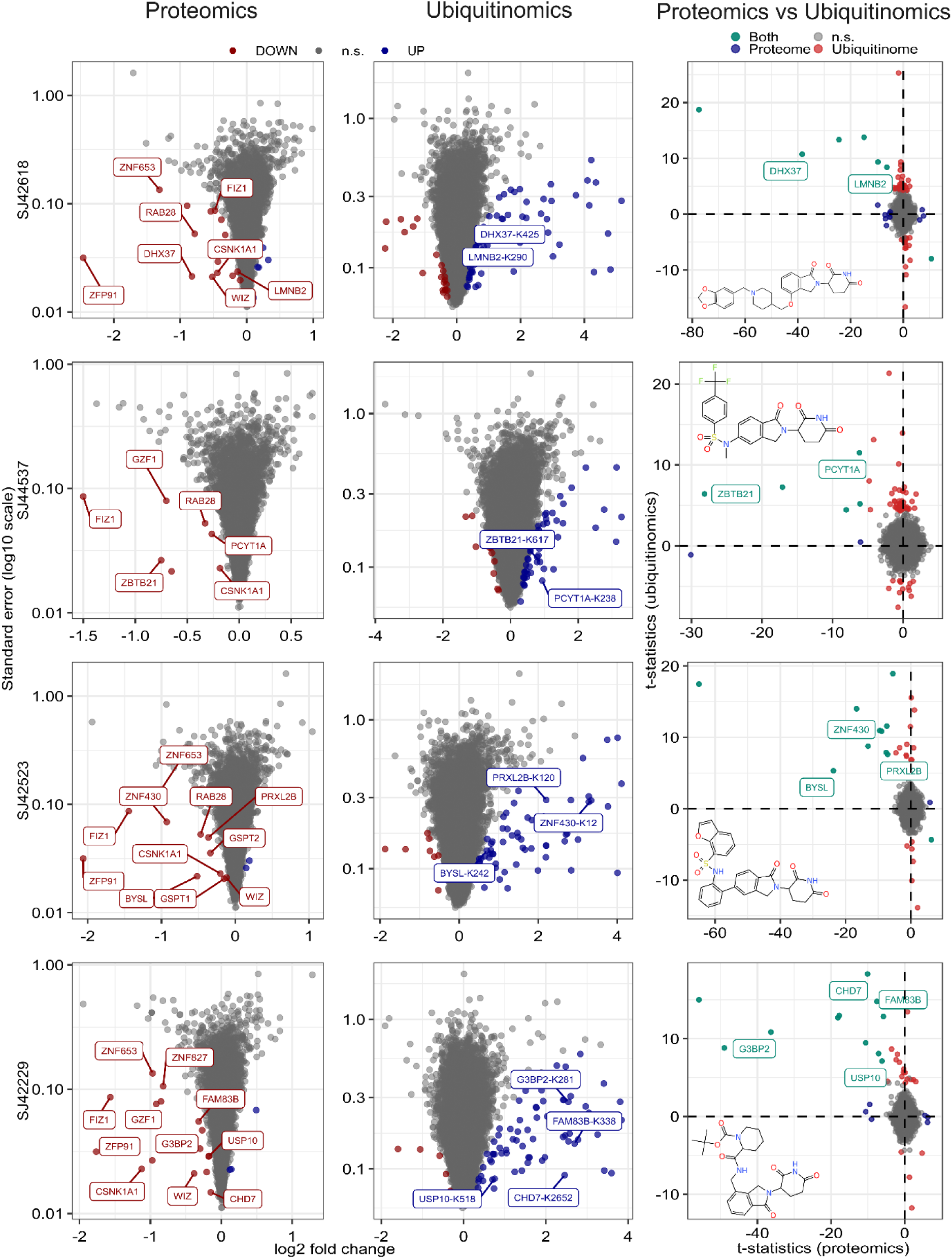
Proteomics and ubiquitinomics of 8 degrader compounds in HEK293-CRBNoe cells. Volcano plots illustrating the proteomes (right panel, 6-hour treatments) and corresponding ubiquitinomes (middle panel, 30-minute treatments) for the indicated MGDs. The x-axis represents the log_2_-transformed fold change, while the y-axis displays the standard error on a log_10_ scale. Significantly upregulated proteins (LIMMA^29^, FDR = 1%) are marked in blue, and significantly downregulated proteins are marked in red. Similarly, significantly ubiquitinated peptides (LIMMA^29^, FDR = 5%) are also highlighted in blue and red, respectively. On the right, a t-statistic comparison plot for each tested compound is presented. Significantly modulated proteins are labeled in blue, ubiquitinated proteins in red, and proteins that are both significantly downregulated and exhibit upregulated ubiquitination sites (or vice versa) are colored in turquoise.

**Figure S12.**
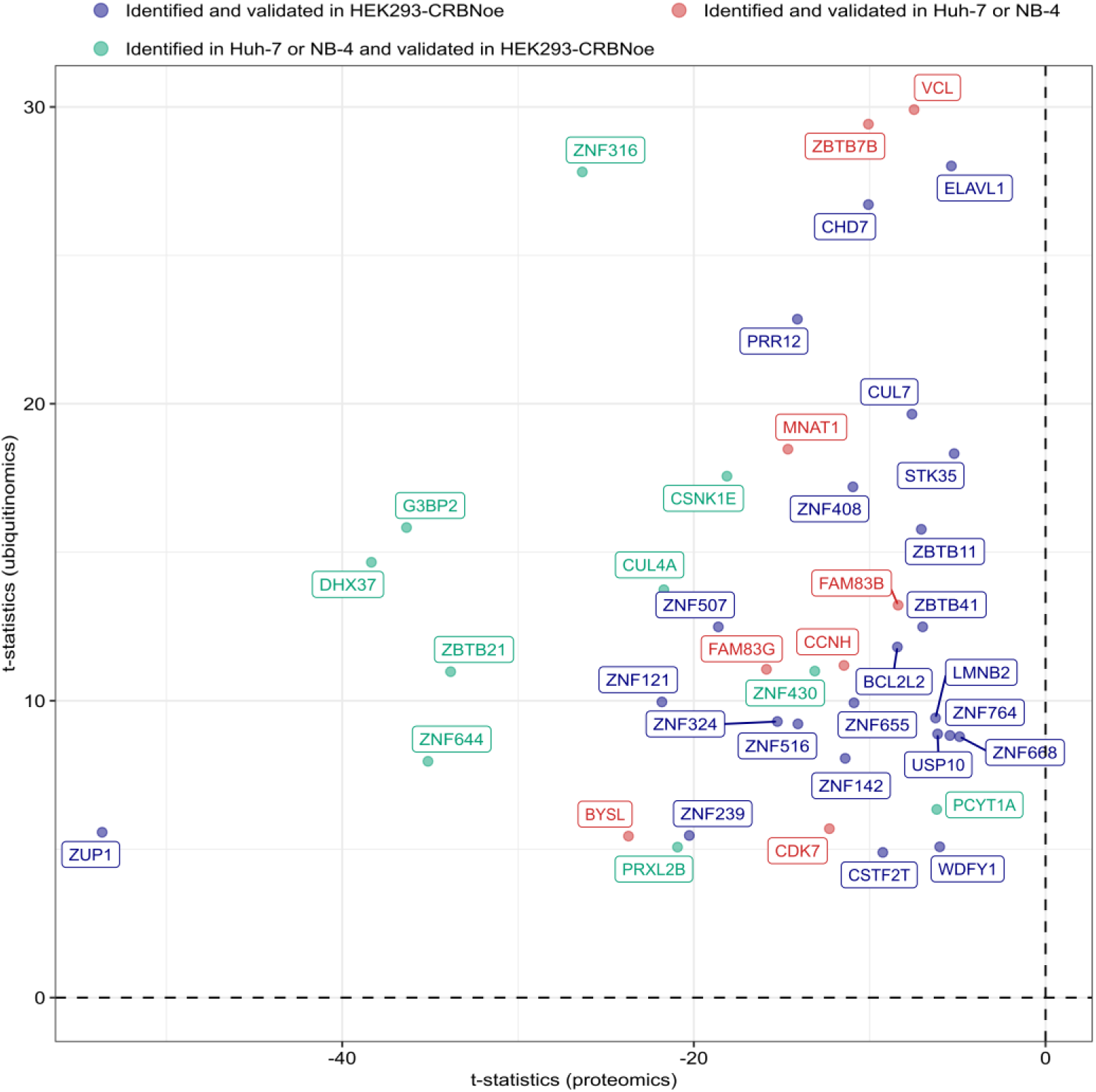
Neosubstrates validated in HEK293-CRBNoe cells. A proteomics versus ubiquitinomics t-statistic comparison plot for each newly validated neosubstrates in HEK293-CRBNoe is presented. Proteins identified and validated through ubiquitinomics specifically in HEK293-CRBNoe cells are shown in blue. Proteins identified and validated in HEK293-CRBNoe, Huh-7, or NB-4 cells are depicted in red. Neosubstrates identified in either NB-4 or Huh-7 cells, but subsequently validated in HEK293-CRBNoe cells, are represented in green.

**Figure S13.**
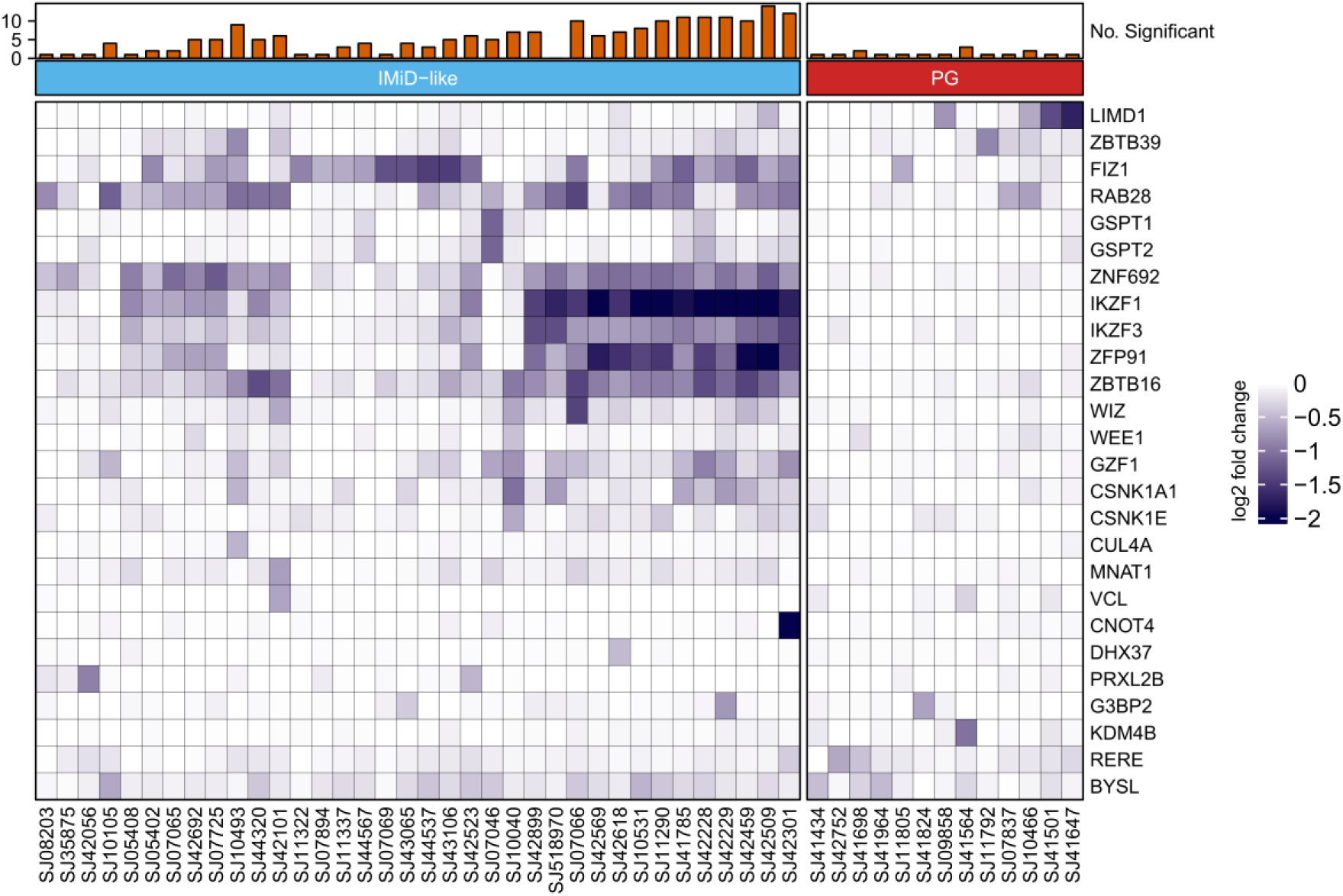
Heap map of selected targets identified in NB-4 cells. Heat map of the log_2_-transformed fold changes of known and selected newly identified neosubstrates in NB-4. The compounds were arranged according to whether they contained a IMiD-like or a PG core structure. The bar chart on top shows the number of significantly downregulated neosubstrates for each compound.

**Figure S14.**
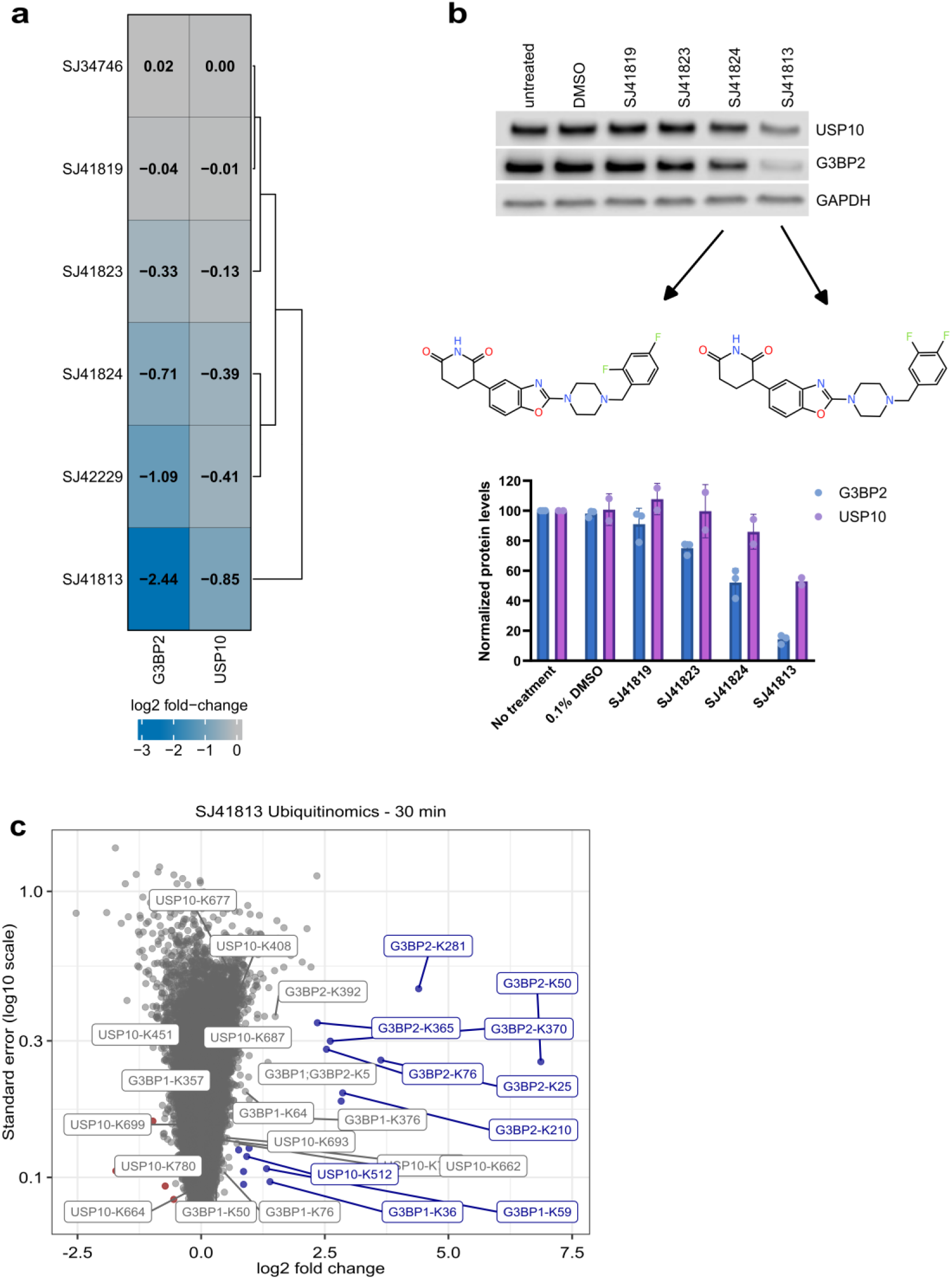
Characterization of G3BP2 degrader compounds. (A) Heat map showing proteomics-derived log_2_ fold-changes for G3BP2 and USP10 following a 24-hour compound treatment with the indicated compounds. (B) Western blot analysis for different analogues of the G3BP2 degrader SJ41824 in SK-N-BE(2)-C cells. Cells were treated with 10 µM of indicated compounds for 24 hours. Depicted blots are representative of two independent experiments. (C) Volcano plot of ubiquitinated peptides upon treatment of HEK293-CRBNoe cells with SJ4813. The x-axis represents the log_2_-transformed fold change, while the y-axis displays the standard error on a log_10_ scale. Significantly up- and downregulated ubiquitinated peptides are marked in blue and red, respectively. Non-significant (n.s.) peptides are colored in grey.

**Figure S15.**
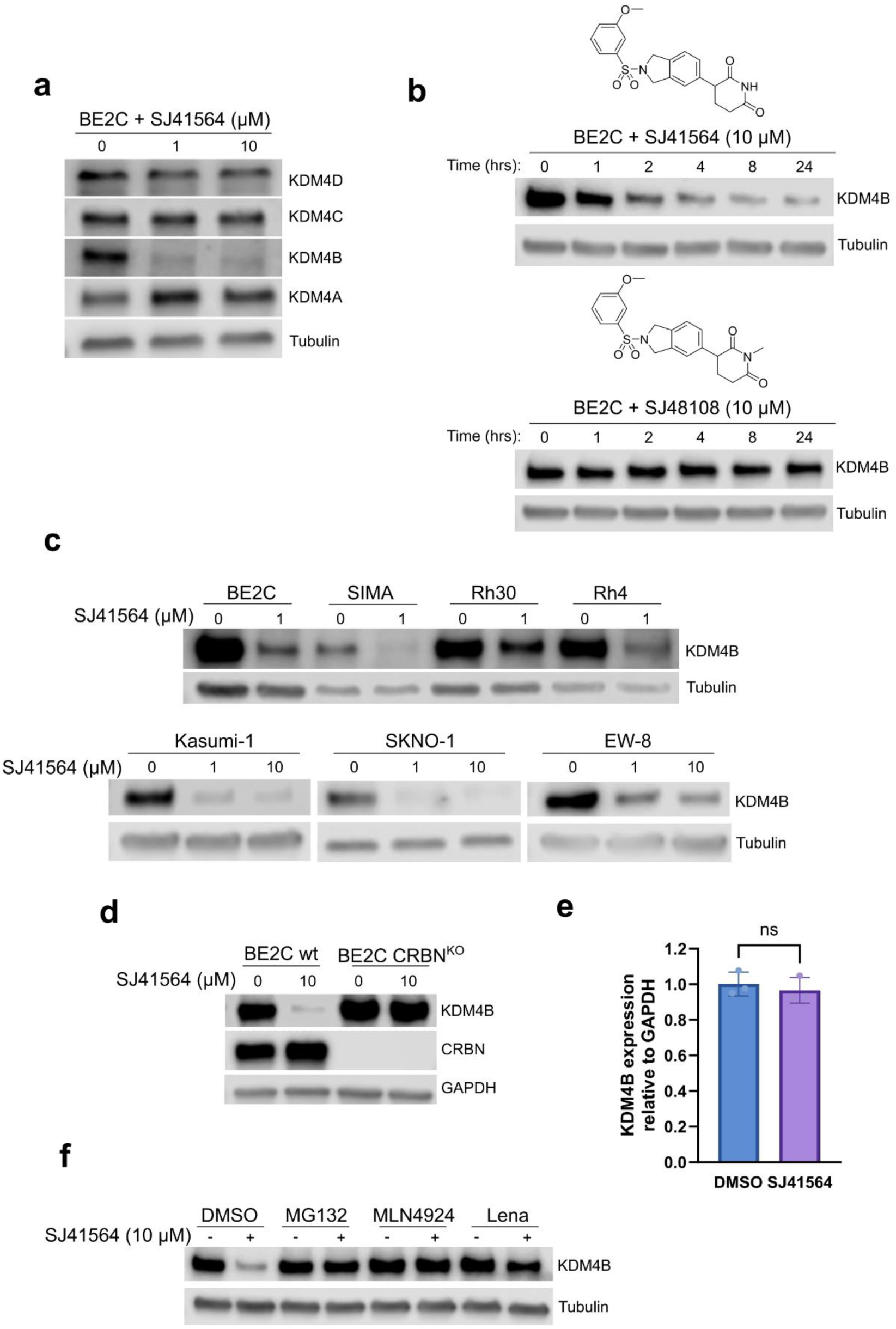
Characterization of the KDM4B degrader SJ41564. (A) Western blot analysis for KDM4 family members upon treatment of SK-N-BE(2)-C cells (BE2C) with DMSO or the degrader SJ41564 for 24 hours (1 and 10 µM). (B) Immunoblots for KDM4B protein after the treatment of SK-N-BE(2)-C cells with increasing concentrations of SJ48108 for 24 h. Depicted blots are representative of three independent experiments. (C) Immunoblots for KDM4B protein after the treatment of SK-N-BE(2)-C cells with 10 µM of SJ41564 over 24 h, and proteins were harvested at 0, 1, 2, 4, 8, and 24 hours after treatment. Depicted blots are representative of three independent experiments. The chemical structure of both compounds is shown. (C) Immunoblots for KDM4B protein after the treatment of neuroblastoma (SK-N-BE(2)-C (BE2C), SIMA), rhabdomyosarcoma (Rh30, Rh4), leukemia (Kasumi-1, SKNO-1), and Ewing sarcoma (EW-8) cells with 1 or 10 µM of SJ41564 for 24 h. (D) Western blot analysis for the indicated proteins of SK-N-BE(2)-C cells (wt and CRBN knockout (CRBN^KO^)) treated with DMSO or 10 µM of SJ41564 for 24 hours. (E) RT-qPCR analysis of KDM4B mRNA level after the treatment of SK-N-BE(2)-C cells with 10 µM of SJ41564 for 24 hours. (F) Immunoblots for KDM4B protein after the treatment of SK-N-BE(2)-C cells with indicated compounds. Cells were pre-treated with MG132 (5 µM), MLN4924 (1 µM), and lenalidomide (20 µM) for 1 h prior to treatment with 10 µM of SJ41564 for 5 hrs.

**Figure S16:**
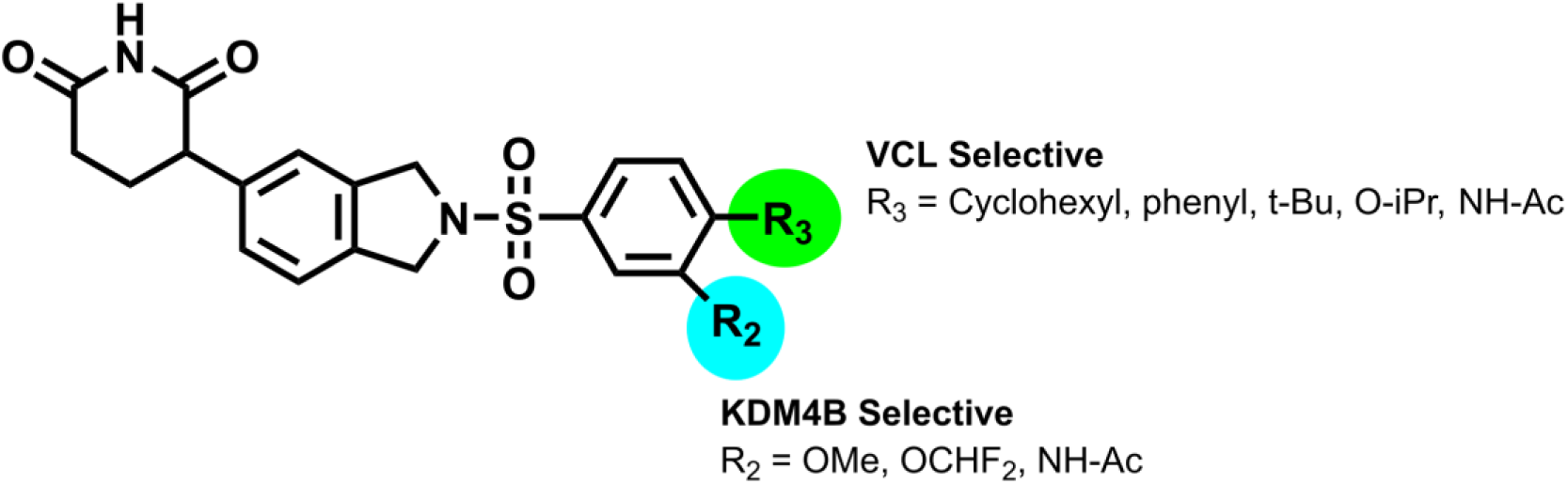
Chemical characterization of KDM4B degrader compounds. Structure-degradation relationship of scaffold illustrating substitution pattern associated with KDM4B and VCL degradation.

**Figure S17:**
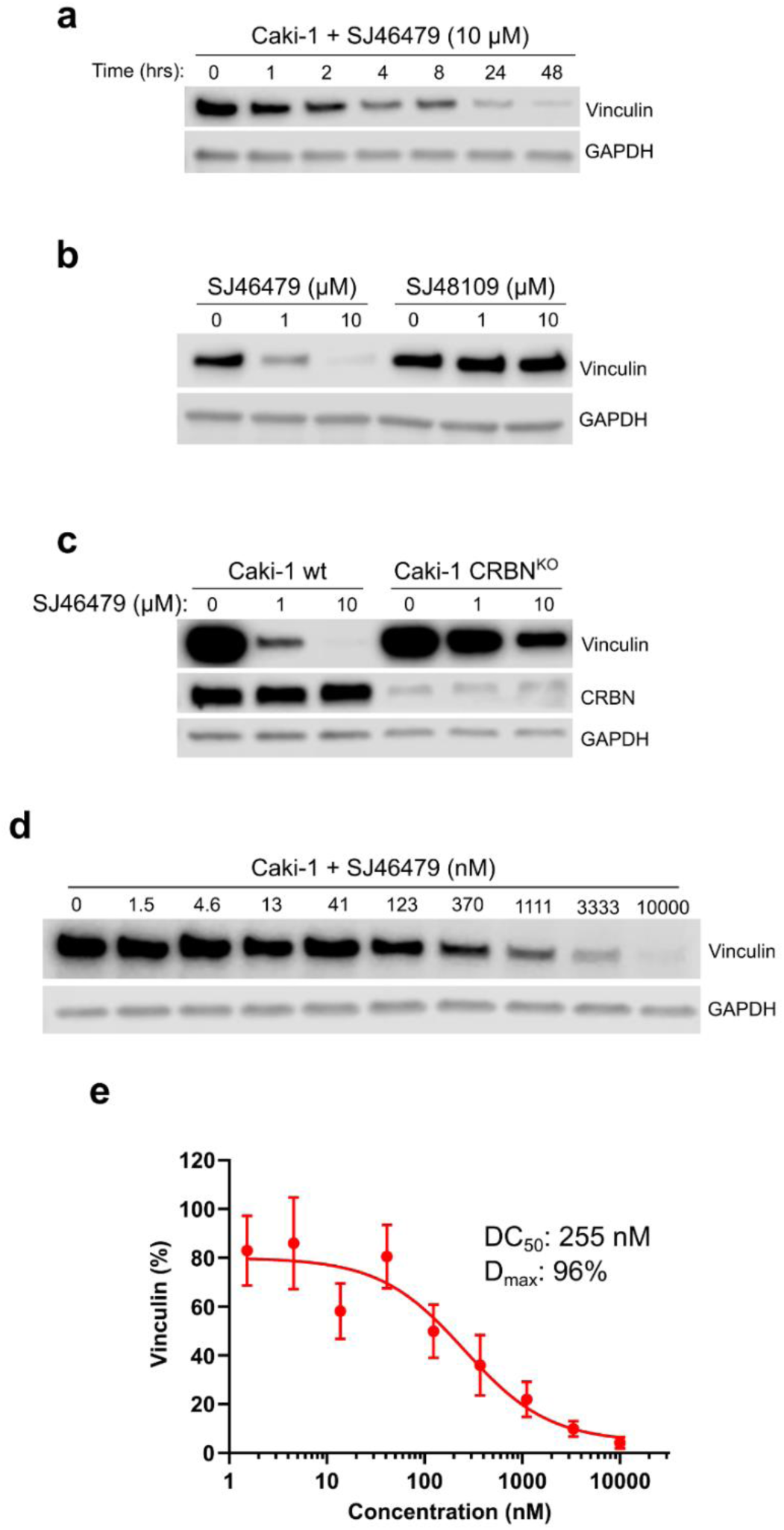
Characterization of the VCL degrader SJ46479. (A) Western blot analysis for vinculin (VCL) following treatment of Caki-1 cells with 10 µM of SJ46479 for the indicated times. (B) Western blot analysis for vinculin (VCL) following treatment of Caki-1 cells with the indicated doses of SJ46479, and the negative control compound SJ48109 for 24 hours. (C) Western blot analysis for vinculin (VCL) in Caki-1 cells (wt or CRBN knockout (CRBN^KO^)) with the indicated doses of SJ46479 for 24 hours. (D) Western blot analysis for vinculin (VCL) following treatment of Caki-1 cells with the indicated doses of SJ46479 for 24 hours. Depicted blots are representative of three independent experiments. (E) Quantification of vinculin (VCL) from the Western blot shown in (D).

## Acknowledgements

Figure 1a was generated with Biorender (Agreement number RM27FBDPFP).

We are grateful for the support of the American Lebanese Syrian Associated Charities (ALSAC) and St. Jude Children’s Research Hospital. This work was also supported in part by the Alex’s Lemonade Stand Foundation Crazy 8 award. Jun Yang was partly supported by American Cancer Society-Research Scholar (130421-RSG-17-071-01-TBG, J.Y.), the National Cancer Institute (1R01CA229739-01 and 1R01CA266600-01A1, J.Y.), Comprehensive Cancer Center core grant CA021765, and the American Lebanese Syrian Associated Charities (ALSAC). The content is solely the authors’ responsibility and does not necessarily represent the official views of the National Institutes of Health. NEOsphere Biotechnologies GmbH gratefully acknowledges funding for this research from the German Federal Ministry of Education and Research (BMBF, grant number 16LW0372). We thank Dr. Jutta Fritz for her support in project managment.

## Author contributions

M.S. designed and performed the experiments, analyzed and interpreted the data, prepared the figures and wrote the manuscript. B.Sch. developed the data analysis pipeline and interpreted the data. B.Sh. analyzed and interpreted the data, edited the manuscript and prepared the figures. U.O. analyzed and interpreted the data. H.D. and Z.R. conceived the study, designed the experiments, analyzed and interpreted the data, and wrote the manuscript. A.H.B., S.M, T.G. and D.B. performed the experiments, co-developed the ubiquitinomics and TurboID assays and analyzed and interpreted the data. V.D. provided software tools essential for raw data processing. G.N. designed the molecular glue library, selected compounds for proteomics-directed medicinal chemistry and edited the manuscript. K.M designed and synthesized compounds. Z.S., J.P., L.Y. synthesized and purified compounds. A.M. performed assay profiling of compounds. Q.W. performed the experiments, analyzed and interpreted the data, edited the manuscript and prepared the figures. Y.J. designed and supervised the study, interpreted the data and was involved in manuscript writing and editing. All authors read and approved the final manuscript.

## Methods

### Reagents

MLN4924, bortezomib, 2-chloroacetamide, tris(2-carboxyethyl)phosphine hydrochloride (TCEP), sodium deoxycholate, Biotin, Na_2_HPO_4_, 3-(N-morpholino)propanesulfonic acid (MOPS), Tris(hydroxymethyl)-aminomethan, trifluoracetic acid, sodium chloride, formic acid and acetonitrile were from Merck. MG132 (Cat# HY-13259) and Lenalidomide (Cat# HY-A0003) were purchased from MedChem Express (MCE). Protease inhibitor mix from ThermoFisher Scientific (A32955). Trypsin from Promega. PTMScan® HS Ubiquitin/SUMO Remnant Motif (K-ε-GG) Kit (#59322) from Cell Signaling Technology.

### Cell culture, drug treatments and cell lysis (Proteomics)

Huh-7, NB-4, HEK293T and HEK293 cells were from Cytion. Huh-7, HEK293T and HEK293s were cultured in DMEM (VWR) supplemented with 10% FCS (Thermo Fisher Scientific). NB-4 were grown in RPMI (VWR) and 10 % FCS. The SK-N-BE(2) (BE(2)-C) cell line was obtained from ATCC. Cells were cultured in RPMI 1640 (Corning, Cat# 10040CM) supplemented with 10% Fetal Bovine Serum (Gibco, Cat# A5256701) and 1% Penicillin/Streptomycin (Gibco, Cat# 15140122). All compounds were dissolved in DMSO, to generate a 1000x stock solution. Compounds were plated in 96-well plates according to a randomized layout using an Opentrons OT-2. Cells were treated with either DMSO or the indicated compounds for the specified amount of time, washed with PBS and harvested using iST lysis buffer (PreOmics). The lysates were heated to 95°C for 10 min while shaking in a Thermomixer (Eppendorf). The rest of the workflow (tryptic digestion and desalting of peptides) was performed according to manufacturer’s instructions.

### Ubiquitinomics sample preparation

Cell treatment and sample preparation was done according to our recently established protocol with a few modifications^28^. In brief, cells were cultured in 6-well plates and treated for 30 min with the specified compounds, followed by lysis with SDC buffer. Protein concentrations were determined using the BCA assay (Merck-Millipore) and the proteins were digested overnight at 37°C using 100:1 protein:trypsin ratio (Promega). After digestion, immunoprecipitation (IP) buffer (50 mM MOPS pH 7.2, 10 mM Na_2_HPO_4_, 50 mM NaCl) was added to the samples together with K-GG antibody-bead conjugate, followed by a 2 h incubation on a rotor wheel. Beads washing and peptide elution was performed according to manufacturer’s instructions. The peptide eluate was desalted using in-house prepared, 200 µl two plug C18 StageTips (3M EMPORE^TM^)^59^.

### Enrichment of biotinylated proteins

After inducing the expression of TurboID-CRBN overnight using 100 ng/ml of doxycycline, the cells were treated with the indicated degrader compounds for 2 hours (in the presence of 50 mM biotin and 0.5 µM bortezomib). Cells were lysed with cold RIPA buffer (1% NP-40, 0.5% SDC, 0.1% SDS, 50 mM Tris-HCl pH=7.5, 150 mM NaCl, freshly supplemented with Protease inhihitor mix) and the lysates were cleared by centrifugation (20,000g, 10 min). 10 µl of MagReSyn® Streptavidin (Resyn Biosciences) was added to the samples followed by 1 hour incubation on a rotor wheel. Beads were washed for 5 times with RIPA buffer and proteins subsequently eluted using UDM (n-Undecyl-Beta-Maltoside, Anatrace) lysis buffer (0.05% UDM, 75 mM Tris-HCl pH 8.5, 40 mM CAA, 10 mM TCEP). The lysates were incubated at 90°C for 10 min and proteins were digested overnight at 37°C using trypsin. The peptides were desalted using in-house prepared, 200 µl two plug C18 StageTips (3M EMPORE^TM^) and resuspended in 0.1% formic acid before MS analysis.

### LC-MS/MS measurements

Peptides were loaded on 35 cm reversed phase columns (75 µm inner diameter, packed in-house with C18-AQ 1.9 µm resin [ReproSil-Pur®, Dr. Maisch GmbH]). The column temperature was maintained at 55°C using a column oven. Either an EASY-nLC 1200 or a Vanquish Neo system (both ThermoFisher) was directly coupled online with the mass spectrometer (timsTOF pro2, HT or Ultra2, Bruker) via a nano-electrospray source. The LC flow rate was 300 nl/min and the complete gradient was 60 minutes (proteomics) or 45 minutes (ubiquitinomics). Data acquisition was done using diaPASEF^18^ or slicePASEF^60^ (for ubiquitinomics) methods.

### Plasmids and cloning

pCMV-FLAG-KDM4B (1-1096), pCMV-FLAGKDM4B (1-130), pCMV-FLAG-KDM4B (130-356), pCMV-FLAG-KDM4B (336-916), pCMV-FLAGKDM4B (336-1096) and pCMV-FLAG-KDM4B (917-1096) were produced as reported previously^61^.

### Generation of stable cell lines

The plasmid vectors for stable cell line generation (CRBN and TurboID-CRBN) were from VectorBuilder. The plasmid vector pLC-Flag-CRBN-P2A-Hygro for generation of stable BE2C cell line overexpressing CRBN was a gift from Eva Gottwein (Addgene plasmid #124303). HEK293 cells with stable gene overexpression were generated by lentiviral-mediated infection of cells according to published protocols^62^. In brief, HEK293T were co-transfected with psPAX2, pMD2.G, pRSV-Rev and a lentiviral construct for CRBN expression under the control of a EF1A promoter. The supernatant was harvested after 48 hours of cell transfection and passed through a 0.45 µm filter. HEK293 cells were infected with the supernatant for 24 hours, followed by selection of transduced cells with puromycin (1 µg/ml). CRBN overexpression was verified by MS-based proteome profiling. HEK293-TurboID-CRBN cells were generated using the PiggyBAC system. Cells were co-transfected with a vector expressing the PiggyBAC transposase and a second vector expressing both TurboID-CRBN (tetracycline-inducible) and rtTA/tTS (Reverse tetracycline responsive transcriptional activator M2/Tetracycline transcriptional silencer), including a puromycin selection marker. Cells stably transfected with the construct were selected with 1 µg/ml puromycin and doxycycline-dependent expression of TurboID-CRBN was verified by mass spectrometry.

### Total RNA extraction, cDNA synthesis, and RT-qPCR

Total RNA was extracted from cells and tumor tissues using RNeasy Mini Kit (Qiagen, Cat # 74106) according to the manufacturer’s instructions. After extraction, 0.5 - 1 µg of total RNA cDNA was used to synthesize cDNA using SuperScript IV First-Strand Synthesis System (Invitrogen, Cat # 18091050) according to the manufacturer’s instructions. RT-qPCR was performed using SYBR Green PCR Master Mix Green (Applied Biosystems, Cat # 4367660) and the 7500 Real-Time PCR System (Applied Biosystems). The relative expression of each gene was normalized to GAPDH mRNA using the ΔΔCT methods. The sequences of primers used are listed below.

KDM4B-F: TCACCAGCCACATCTACCAG

KDM4B-R: GATGTCCCCACGCTTCAC

GAPDH-F: AACGGGAAGCTTGTCATCAATGGAAA

GAPDH-R: GCATCAGCAGAGGGGGCAGAG

### Antibodies and Western blot

Cells were lysed on ice using a lysis buffer (0.1 M Tris-HCl, pH 6.8, 200 mM dithiothreitol, 0.01% bromophenol blue, 4% sodium dodecyl sulfate and 20% glycerol). Samples were sonicated at 4 °C for 10 s, and then boiled at 95 °C for 10 min. Cell lysates were separated on 4–15% precast polyacrylamide gel (BioRAD, Cat # 4568086) and transferred to PVDF membranes (BioRad, Cat # 1704272) pre-activated with methanol. Membranes were blocked in a solution of 5% milk in PBST buffer (0.1% TWEEN in PBS) and incubated for 1 hour at room temperature and incubated overnight with primary antibodies at 4 °C under gentle horizontal shaking. The next day, membranes were washed with PBST buffer and incubated with anti-mouse or anti-rabbit HRP-conjugated secondary antibodies (1:5,000). Membranes were washed with PBST buffer, incubated with SuperSignal West Pico PLUS Chemiluminescent Substrate (ThermoFisher, Cat # 34580) and developed using an Odyssey Fc Imaging System (LI-COR Corp.). Antibodies and working concentrations are provided in the table below.

| Antibody | Source | Identifier | Concentration |
| --- | --- | --- | --- |
| GAPDH (14C10) | Cell Signaling Technology | Cat# 3683S<br>RRID: AB_1642205 | 1:2000 |
| CRBN (D8H3S) | Cell Signaling Technology | Cat# 71810S<br>RRID: AB_2799810 | 1:1000 |
| G3BP2 | Bethyl Laboratories | Cat# A302-040A<br>RRID: AB_1576545 | 1:1000 |
| USP10 (D7A5) | Cell Signaling Technology | Cat# 8501S<br>RRID: AB_10949976 | 1:1000 |
| KDM4A | ABclonal | Cat# A9267<br>RRID: AB_2863701 | 1:1000 |
| KDM4B | Bethyl Laboratories | Cat# A301-478A<br>RRID: AB_999594 | 1:1000 |
| KDM4C | ABclonal | Cat# A8485<br>RRID: AB_2770070 | 1:1000 |
| KDM4D | ProteinTech | Cat# 22591-1-AP<br>RRID: AB_2879130 | 1:1000 |
| Flag M2 (F1804) | SIGMA | Cat# F1804<br>RRID: AB_262044 | 1:1000 |
| α-Tubulin (6A204) | Santa Cruz Biotechnology | Cat# sc-69969<br>RRID: AB_1118882 | 1:1000 |
| HSP90$\alpha/\beta$ (F-8) | Santa Cruz Biotechnology | Cat# sc13119<br>RRID: AB_675659 | 1:1000 |

### Raw data processing

MS raw files were analyzed using DIA-NN^10^. Proteomics raw files were processed with v1.8.2beta9 or 1.9.1 (chemical analog testing) and ubiquitinomics raw data was processed with v1.8.2beta27. Reviewed UniProt entries (human, SwissProt 10-2022 [9606]) were used as protein sequence database for DIA-NN searches. One missed cleavage, a maximum of one variable modification (Oxidation of methionines) and N-terminal excision of methionine were allowed. Carbamidomethylation of cysteines was set as fixed modification and K-GG (UniMod: 121) was added in case of ubiquitinomics. All data processings were carried out using library-free analysis mode in DIA-NN. --tims-scan was added as additional command in case of ubiquitinomics.

### Statistical data analysis

DIA-NN outputs were further processed with R. Peptide precursor quantications with missing values in more than 50% of samples, or <33% of the DMSO-treated samples (for proteomics, <25% of compound-treated samples in case of ubiquitinomics) were discarded. Protein abundances were calculated using both precursor and fragment ion intensities; K-GG peptide abundances were calculated based on precursor ion intensities levels using the MaxLFQ algorithm^63^, as implemented in the DIA-NN R package (https://github.com/vdemichev/diann-rpackage/). Missing for cases with complete missingness were replaced by low-quality precursors (i.e., q-value > 0.01). K-GG peptide to site mapping was done using reviewed entries of the human UniProt database (SwissProt, release 10-2022). The protein (or peptide) intensities were normalized by median scaling and corrected for variance drift over time (if present) using the principal components (derived from principal component analysis (PCA)) belonging to DMSO samples. Subsequently, protein (or peptide) intensities were subjected to statistical testing with variance and log fold-change moderation using LIMMA^29^. p-values corrected for multiple testing were used to assess significance for proteomics (q-value < 0.01) and ubiquitinomics (q-value < 0.05). For comparing proteome and ubiquitinome data, identifications were mapped at the gene level.

